# Probabilistic Co-Control in Brain-Computer Interfaces: Uncertainty as a Control Signal in Brain-to-Text Decoding

**DOI:** 10.64898/2026.04.02.715749

**Authors:** Jingya Huang, Sowmya Manojna Narasimha, Aashish N. Patel, Ram Dyuthi Sristi, Gal Mishne, Vikash Gilja

## Abstract

Neural decoders serve as probabilistic interfaces in co-control brain-to-text BCIs, where predicted uncertainty shapes hypothesis generation and language model integration, enabling decisions to be made safely under uncertainty. However, it remains unclear whether these decoders produce reliable and informative uncertainty, or how training objectives shape these properties. This work characterizes and improves uncertainty representations in brain-to-text decoding. We extend two metrics, calibration error (ECE) and resolution (RES), to evaluate sequential probabilistic predictions from frame-level phoneme estimates to word-level hypotheses, quantifying the reliability and informativeness of model uncertainty. Using this framework, we analyze neural decoders trained with connectionist temporal classification (CTC). To isolate the causal role of uncertainty independent of accuracy, we manipulate predicted probability distributions while holding predicted sequences fixed. Motivated by the observed failures, we further examine the role of the training objective and propose a two-stage cross-entropy (CE) formulation that decouples alignment inference from classification. We show that widely used CTC-trained neural decoders in brain-to-text BCIs produce systematically over-confident predictions, with high confidence persisting even when predictions are incorrect. Controlled manipulations of the prediction reveal that improved ECE and RES enhance hypothesis generation and language-model integration by promoting diverse alternatives and more effective re-ranking of hypotheses aligned with user intent. Mechanistically, CTC relies on over-confident predictions to resolve alignment ambiguity. Replacing CTC with CE loss yields significantly more reliable and informative probabilistic predictions without degrading decoding accuracy. Uncertainty emerges as a system-level design variable in brain-to-text interfaces. Calibrated uncertainty from neural decoders enables effective integration with independently trained language models and reliable error detection. This work reframes uncertainty from a passive output into an active control signal, identifies key components and evaluation criteria for probabilistic co-control, and outlines a pathway toward next-generation BCIs that supports increasingly complex interactions with the world.

## 1 Introduction

Brain–computer interfaces (BCIs) aim to restore communication and movement [1, 2] in a manner that is effective, safe, and minimizes cognitive burden [3, 4], ultimately restoring independence for individuals with paralysis. Decoded neural signals may directly drive assistive devices that facilitate daily activities through communication interfaces [5–12], software applications [13, 14], wheelchair or robotic limbs [15–23]. As a result, deployed BCIs are often safety-critical, as misinterpreted user intent can trigger conflicts or irreversible actions [3, 23, 24]. In this setting, average predictive accuracy alone is insufficient, though it is the performance measure most commonly reported. Uncertainty estimates provide an informative signal when a system’s outputs should be treated with caution or deferred for intervention as established in safety-critical domains (e.g., robotics and medical diagnosis [25–30]). Reliable and informative uncertainty estimates are therefore a fundamental requirement for practical BCI deployment.

The need for high-quality uncertainty estimates is amplified in modern BCI architectures that employ co-control [31, 32], where neural decoders and behavior models (e.g., language models, motion planners) jointly determine control outputs. As BCIs scale to more complex behaviors, decoding must resolve a large space of possible outputs, typically requiring substantial paired neural–behavior data. However, such data are difficult to collect, especially for users who require assistance, and are often limited in both quantity and precision. Brain-to-text BCIs exemplify this challenge [11,12,33–39], where participants cannot produce overt speech, handwriting, or typing. Instead, intended text is inferred through experimental protocols. As a result, supervision is weak: it lacks fine-grained temporal alignment between neural activity and behavior and obscures trial-to-trial variability in the underlying behavior. Co-control mitigates these limitations by leveraging independently trained behavior models that encode statistical regularities over behavior. These models act as priors over the space of possible outputs, constraining decoding and thereby reducing reliance on large amounts of paired neural–behavior data while improving performance. Co-control also enables the extensible integration of additional components, supporting scalable and modular BCI design (Fig. 1).

**Figure 1:**
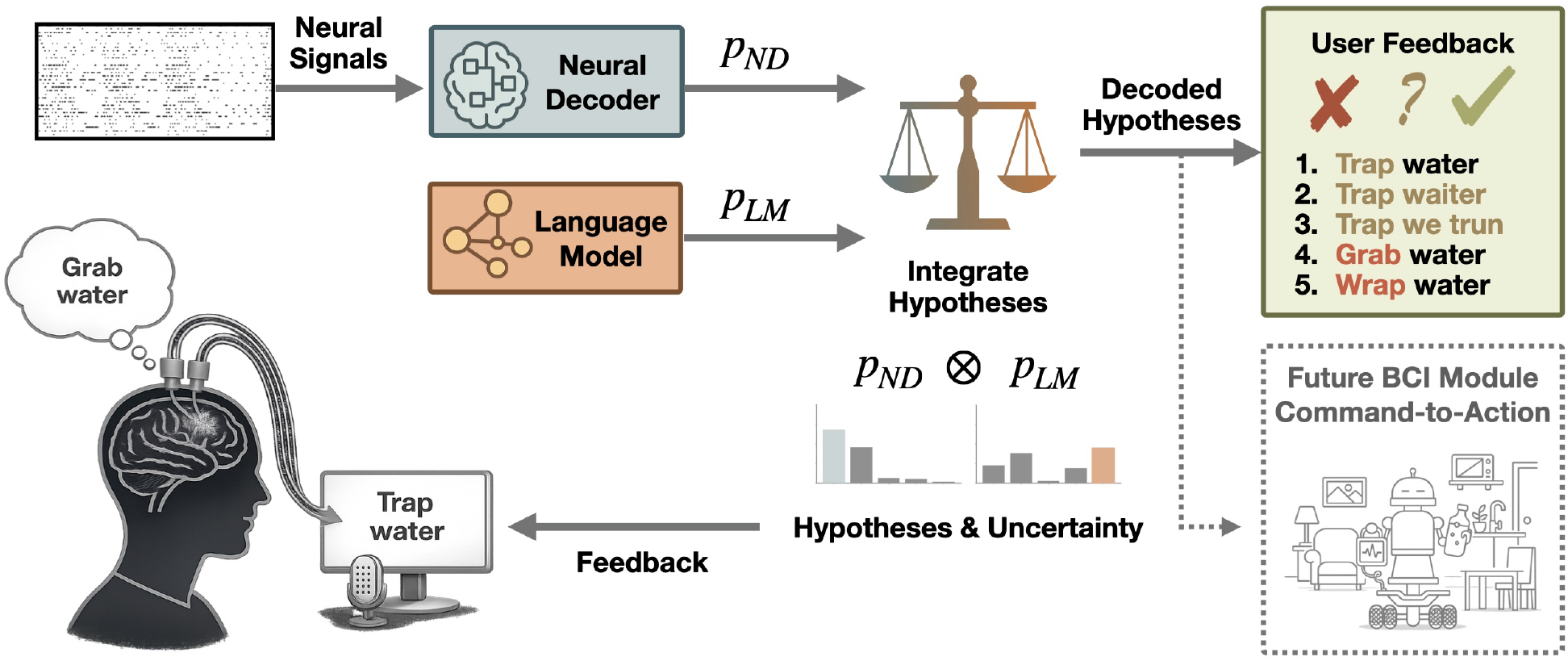
Co-control Brain-Computer Interface (BCI) prototyped with the brain-to-text system. Schematic of a co-control BCI in which decoded intent and estimated uncertainty propagate through inference, integration, and action layers to guide system behavior. A neural decoder maps intracortical signals to a predictive distribution *p*_ND_ over candidate text sequences. These distributions are combined with language model priors *p*_LM_ via ranked arbitration (*p*_ND_ ⊗ *p*_LM_) to produce the decoded hypotheses, with performance governed by the uncertainty encoded in *p*_ND_. Beyond decoding, uncertainty serves as an interface for interaction: it highlights likely errors for user feedback, supports efficient correction, and enables safety-aware downstream control by gating actions based on uncertainty. This framework illustrates how uncertainty functions as a central mechanism for reliable probabilistic co-control in BCI systems, demonstrated here in the context of discrete brain-to-text decoding.

Although brain-to-text BCIs are often framed as directly mapping neural activity to text, existing systems are more accurately understood as inherently operating under co-control. Co-control refers to the joint construction of control signals in a shared hypothesis space, where multiple modules contribute by weighting candidate hypotheses according to their estimated uncertainty. In practice, this is realized through pretrained language models [40–45] correcting, and refining neural decoder outputs [11, 12, 35–39, 46–49]. Similar co-control paradigms have emerged in BCI-enabled high-degree-of-freedom reach-and-grasp robotic control tasks [22, 50, 51]. In these modular pipelines, neural decoders operate as components within a broader system in which downstream models impose learned behavioral structure and constraints. Although such integration enables substantial performance gains, including languagemodel-guided error correction in brain-to-text decoding [11, 12, 35–39, 46–49, 52, 53] and obstacle-aware motion planning in brain-to-grasp control [22, 50, 51], it also amplifies the importance of how uncertainty is represented and communicated across system components. Miscalibrated uncertainty can lead to failures at both extremes: excessive assistance may override user intent and hinder goal achievement, while insufficient assistance can result in poor control and uncorrected errors, ultimately leading to misalignment between behavior-model assistance and user intent that impacts usability, safety, and user trust [28, 31, 51, 54–56].

Effective co-control in systems like brain-computer interfaces relies on an uncertainty-governed division of labor, much like a collaborative human transcription team. In this analogy, the primary decoder, whether a human transcriber or a BCI neural decoder, serves as the interface that translates raw signals into initial commands, ideally flagging segments of ambiguity or low confidence. A downstream “editor”, such as a human proofreader or a BCI language model, refines these outputs by focusing on flagged regions and using contextual consistency to infer likely corrections without direct access to the original signal (audio or neural activity). Crucially, the success of this collaboration hinges on how accurately uncertainty is communicated between these stages: the editor’s ability to intervene is fundamentally limited if the uncertainty conveyed by the transcriber or neural decoder is misaligned with the underlying signal variability. In the transcription analogy, this corresponds to expressing high confidence despite degraded audio (e.g., noise or overlapping speech), or low confidence when the signal is clear, both of which misguide the editor’s corrections. In the BCI setting, signal variability reflects ambiguity in neural activity, while uncertainty should indicate how reliably the decoder maps that activity to intended phonemes or words. When this alignment fails, downstream modules cannot distinguish between reliable and unreliable predictions, undermining effective co-control. Thus, uncertainty is not a secondary diagnostic, but the central signal that governs arbitration between independently trained components, ensuring that the final output reflects the user’s intent rather than the bias of any single module.

In brain-to-text BCIs, this interaction occurs through probabilistic outputs rather than discrete predictions, allowing downstream modules to reason about uncertainty. Neural decoders produce a distribution over text tokens across time, which is processed by pretrained language models to generate and re-rank candidate sentences [40–45] (Fig. 1). Ideally, elevated uncertainty in the neural decoder output would signal potential errors, expand the set of candidate hypotheses explored during sequential decoding, and shift control toward the language model for correction. However, in practice, the quality of this uncertainty signal is rarely examined explicitly. Most brain-to-text systems are evaluated primarily using end-to-end metrics, such as error rates (e.g., word, phoneme, or character error rate; WER, PER, CER), communication rate (e.g., words or characters per minute; WPM, CPM), or semantic relevance (e.g., BLEU, BERTScore) [11,12,35–39,46–49,52,57–62], obscuring how uncertainty is represented, interpreted, and acted upon within the co-controlled architecture. As a result, neural decoders are often optimized to be “decisive” at the expense of being “informative”. In practice, this can lead to overconfident predictions that disrupt uncertainty-governed co-control, impairing appropriate control allocation under ambiguity and promoting overcorrection that obscures the user’s intended output. This raises several fundamental questions: Does a more accurate neural decoder necessarily lead to better integrated system behavior under co-control? What determines the quality of uncertainty estimates in its predictions? And can uncertainty itself be systematically quantified and can its quality be improved to enable more principled and reliable co-control?

The challenge of obtaining reliable and informative uncertainty estimates in classification is not unique to BCIs. In image classification, deep neural networks can achieve high predictive accuracy while producing poorly calibrated uncertainty estimates [63–66]. This gap between accuracy and reliability has motivated extensive work on uncertainty quantification, including post-hoc calibration methods [64–66], conformal prediction [67,68], Bayesian neural networks [69], and ensemble approaches [70,71]. However, most of these methods are developed for static, instance-level prediction. Extending uncertainty estimation to sequential, structured decoding, where probabilities propagate across time and interact with search and downstream models, is substantially more challenging.

Brain–computer interfaces represent a particularly demanding instance of this broader problem. Neural decoding is sequential and structured, and uncertainty estimates must be reliable to preserve user intent as probabilistic predictions propagate through intermediate inference stages. In this work, we use brainto-text decoding as a prototype for probabilistic co-control, offering a principled framework to study how neural decoder predictions of user intent are combined with complex behavioral priors, such as language models, through probabilistic interfaces. First, we introduce a principled evaluation framework to quantify the reliability and informativeness of probabilistic predictions across multiple temporal scales by leveraging calibration error and resolution, which is defined as the ability of predicted probabilities to distinguish correct from incorrect outcomes, grounded in proper scoring rule decompositions [72]. Second, we show that widely used neural decoders trained with the connectionist temporal classification (CTC) loss produce systematically over-confident predictions. Third, through oracle interventions, we demonstrate that improved calibration and resolution for uncertainty estimates, substantially increase hypothesis diversity and enhance downstream integration with language models. Finally, we provide mechanistic insight by showing that CTC implicitly relies on over-confident predictions to resolve alignment ambiguity. By replacing this objective with a two-stage CTC and cross-entropy (CE) formulation that first stabilizes alignments and then focuses on classification, we obtain neural decoders with significantly more reliable and informative probabilistic predictions, enabling more principled co-control.

## 2 Methods

### 2.1 Brain-to-Text BCI and Dataset

Brain-to-text BCIs decode neural activity during attempted speech to generate text for individuals with paralysis. We analyze publicly available data from participants T12 [12] and T15 [35], recorded during the delayed sentence production task in which participants attempted to speak prompted sentences. Neural activity and cued sentences form paired input–label data for training.

Neural data were recorded from two 96-channel Utah arrays implanted in ventral motor cortical speech regions (T12: 128 channels in areas 44 and 6v across 19 days [12]; T15: 256 channels in area 6v, 4, and 55b [35] across 45 days). We use the preprocessed signals provided with the datasets without modification. These include threshold crossings (−4.5×RMS) and spike-band power, binned at 20 ms, smoothed, and normalized to form input features **X**_1:*T*_ ∈ ℝ^*T*×*D*^. Models were trained and evaluated on data pooled across recording days unless otherwise specified.

As intelligible speech or articulatory kinematics cannot be recorded from these participants, supervision is derived from task prompts. Each sentence is converted into a phoneme sequence **y**_1:*N*_ = (*s*_1_^′^, …, *s*^′^*N*), *s*^′^*n* ∈ 𝒱, while the underlying framewise phoneme sequence ***π***_1:*T*_ = (*s*_1_, …, *s*_*T*_), *s*_*t*_ ∈ 𝒱 is treated as latent. The sentence is obtained by collapsing repeated phoneme states, **y**_1:*N*_ = C(***π***_1:*T*_), abstracting away the unknown onset and duration of phonemes during speech production.

Brain-to-text decoding combines neural decoder likelihoods with language prior during inference. We interpret this process as a form of co-control, in which neural and language models jointly contribute to inference over a shared hypothesis space, with their influence determined by uncertainty. In this work, we analyze the probabilistic formulation of this system, viewing neural decoder predictions and their associated uncertainty as control signals within the co-control framework. The following sections formalize the neural decoding problem, introduce neural decoder variants that predict phoneme distributions from neural activity, and analyze how uncertainty influences decoding.

### 2.2 Sequence Neural Decoder (ND)

A brain-to-text neural decoder (ND) is a probabilistic sequence model that maps neural activity **X**_1:*T*_ to a sequence of predictive distributions over tokens *s*_*t*_ ∈ 𝒱 (e.g., phonemes [12, 35], characters [11], gestures [36]) at each frame *t* ∈ {1, …, *T* }. Each predictive distribution *p*_ND_(*s*_*t*_ | **X**_1:*t*_) defines a time-varying distribution over token predictions and can be interpreted as uncertainty (Fig. 2).

**Figure 2:**
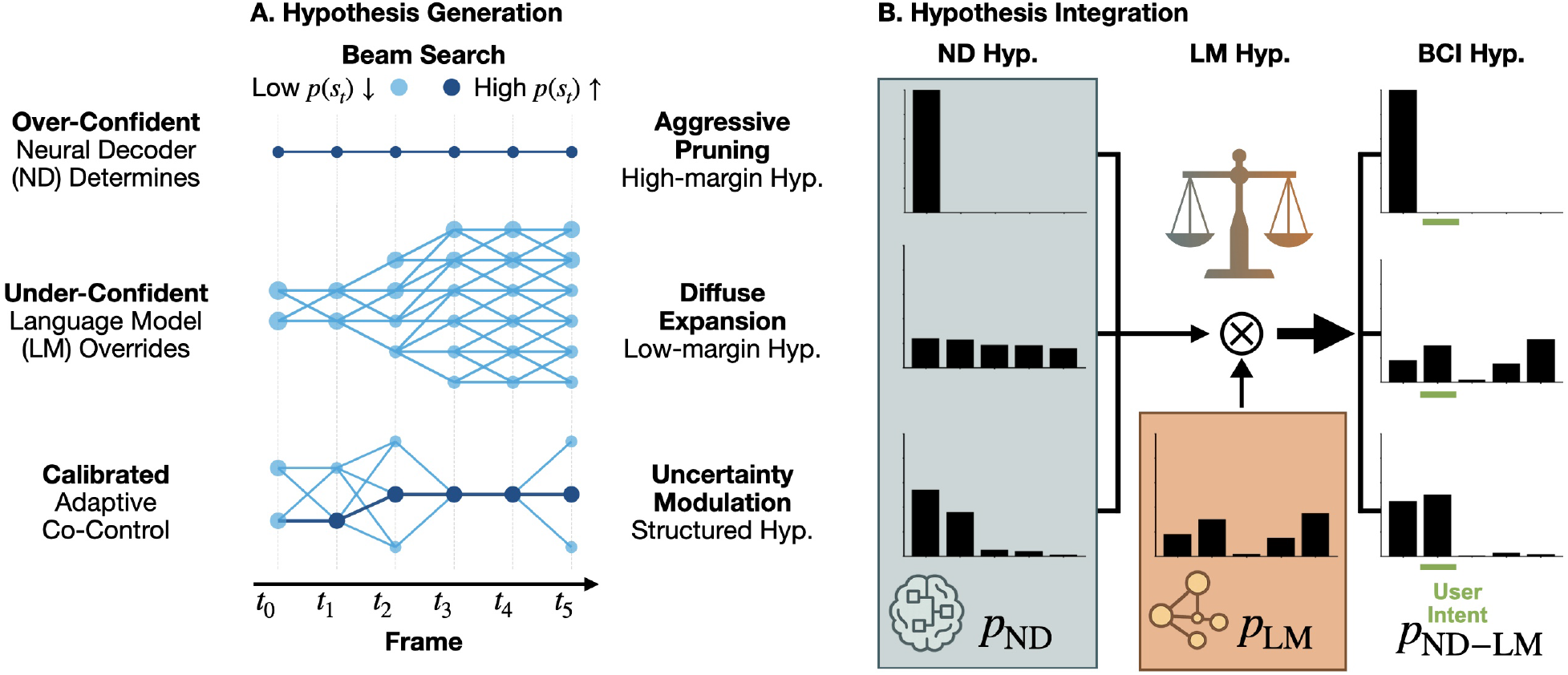
Neural decoder (ND) uncertainty governs hypothesis generation and Language model (LM) integration. Schematic illustrating how ND uncertainty shapes sequence probabilities, which determine (A) the diversity and separation of candidate hypotheses during generation, and (B) how these hypotheses are ranked and integrated with the language model. Three characteristic regimes of *p*_ND_ are shown: over-confident, under-confident, and calibrated. A calibrated *p*_ND_ supports effective and adaptive co-control.

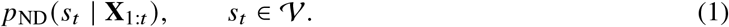

Unlike generic sequence-to-sequence models that learn unconstrained alignments between input **X**_1:*T*_ and output **y**_1:*N*_ sequences, neural decoding operates under a structured but latent temporal alignment, where neural activity and movement production unfold jointly and monotonically over time.

We model decoding as inference over the latent alignment path ***π***_1:*T*_, with model prediction:

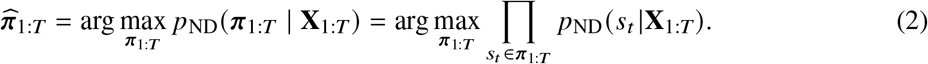

The alignment path encodes token identity and duration, with repeated tokens indicating sustained articulation. Applying the collapse operator C(·) yields the sequence-level hypothesis 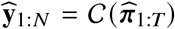, corresponding to the model’s estimate of the intended sequence **y**_1:*N*_.

Supervision for neural decoder training is provided by the intended sentence 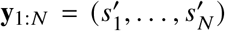). Each sentence (i.e., the prompt-derived phoneme sequence) corresponds to multiple possible alignment paths, defined by the alignment set 𝒜 (**y**_1:*N*_) = {***π***_1:*T*_ : C(***π***_1:*T*_) = **y**_1:*N*_ }

This formulation separates modeling assumptions, framewise factorization, and the collapse mapping from the training objective. Supervision may be imposed either at the sequence level by maximizing the marginal likelihood over alignments with connectionist temporal classification (CTC) loss, or at the frame level using inferred alignments with cross-entropy (CE) loss. The two approaches differ in how supervision is applied to the underlying probabilistic sequence model.

### 2.3 CTC Neural Decoder

Neural decoders are typically trained with sequence-level supervision using the connectionist temporal classification (CTC) objective [11, 12, 35–39, 46–49, 53]. CTC [73] handles latent alignment by maximizing the sequence likelihood via marginalization over all alignments ***π***_1:*T*_ ∈ 𝒜 (**y**_1:*N*_).

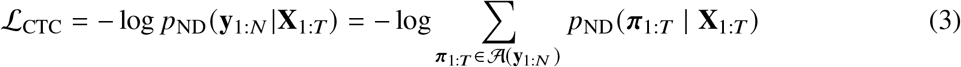

Through this marginalization, CTC jointly performs alignment inference and token classification. This coupling induces “peaky alignment,” where probability mass concentrates on a small number of paths, as widely observed in sequence labeling tasks using CTC loss [74, 75].

### 2.4 CE Neural Decoder

To disentangle alignment from token classification, we use a two-stage framework where CTC-derived framewise alignments are fixed as supervision for a cross-entropy (CE) decoder, with the goal of allowing the predictive distribution to better reflect classification uncertainty.

The most probable CTC alignment 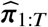 is unconstrained and may collapse to a sequence inconsistent with the ground truth, i.e., 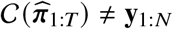. We therefore compute a forced alignment [76]:

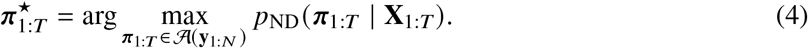

Using 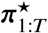 as supervision, the CE decoder is trained with a framewise cross-entropy objective to align the predictive distribution with 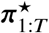.

#### Reinterpreting CTC as iterative CE training

CTC can be viewed as an iterative CE fitting in which predictive distribution conditioned on the target sequence act as soft training targets [77]. At each iteration, the CTC gradient is equivalent to that of a CE loss with soft targets 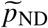:

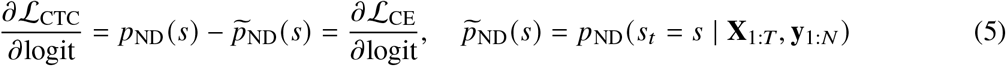

Since these targets 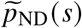 depend on the model parameters, CTC training iteratively fits both alignment and token classification. This coupling provides a mechanism for the emergence of peaky alignments, as the moving targets bias optimization toward a small number of high-probability alignments. In contrast, our approach uses the same final alignment 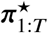 but fixes it as the classification target during CE training. This leads to a different optimization trajectory even when the inferred alignment is identical. If the forced alignment is accurate, the decoder may learn token probabilities that more faithfully reflect classification uncertainty rather than alignment ambiguity.

### 2.5 Fusing Neural Decoder Predictions (CTC ⊗ CE, CTC ⊗ CTC)

To interpolate between alignment-driven and classification-driven regimes, we combine the predictive distributions *p*_CTC_ and *p*_CE_ from CTCand CE-trained models, respectively, using product-of-experts (PoE) [78]. The combined predictive distribution *p*_CTC⊗CE_ is defined as:

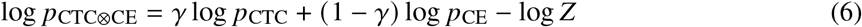

where *Z* is the normalization constant. This formulation defines a continuous family of decoders spanning classification-driven CE (*γ* = 0) and alignment-driven CTC (*γ* = 1). Similarly, we define CTC ⊗ CTC by combining two independently trained CTC models using the same formulation, serving as a control for fusion effects independent of training objective.

### 2.6 Brain-to-Text Decoding Algorithm

Brain-to-text decoding follows a multi-stage probabilistic inference pipeline [11, 12, 35–39, 46–49, 53]. The system first performs framewise neural decoding with ND to estimate phoneme predictive distributions over time, *p*_ND_ (Eq. 1), which are then integrated into sequence probabilities during hypothesis generation and subsequently integrated with a language model (LM). Candidate hypotheses are then identified via lattice-based search over sequence probabilities or fused scores (Fig. 2).

The probability of sequence **y**_1:*N*_ is given by the marginal distribution over all alignments 𝒜 (**y**_1:*N*_):

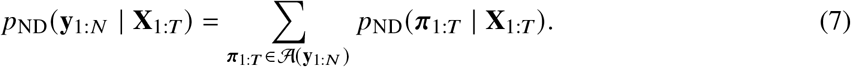

The LM is integrated with ND in log-probability space, with a weight *β* controlling its contribution:

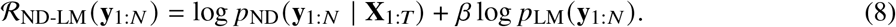

Beam search selects the top-*K* hypotheses 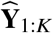 based on *p*_ND_(**y**_1:*N*_ | **X**_1:*T*_), corresponding to sequences generated by the neural decoder alone, or based on the fused score R_ND-LM_(**y**_1:*N*_) over a lattice constructed from a lexicon graph and an n-gram language model using a weighted finite-state transducer (WFST) [41, 42]. This produces top-*K* hypotheses 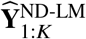 that combine user intent inferred by the neural decoder with linguistic priors from the language model:

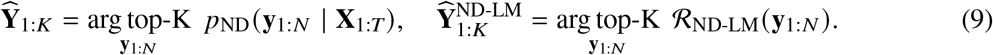

Neural decoder and language model integration operate at the word level, where words define the shared hypothesis space for co-control. Sequence scores decompose additively over word units.

### 2.7 Evaluating Probabilistic Inference at Multiple Timescales

Predictive distributions *p*_ND_ (Eq. 1) provide richer information than point estimates such as the sequence hypothesis 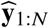 or alignment 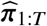 (Eq. 2). While error metrics assess correctness, proper scoring rules evaluate how well predicted probabilities reflect outcome likelihoods. Effective probabilistic predictions require both calibration and resolution, measuring alignment with empirical correctness and the ability to distinguish outcomes, respectively. In co-control BCIs (i.e. brain-to-text), these uncertainty estimates act as control signals guiding hypothesis generation, behavior model integration, and user feedback (Fig. 1), and their evaluation across timescales reveal how uncertainty propagates from frame-level predictions to structured outputs such as words (Fig. 2).

For a probabilistic predictor 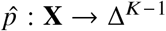 producing 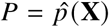, the Brier score 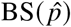 [65] is the proper scoring rule which measures the discrepancy between the predictive distribution 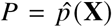 and the realized one-hot outcome *Y*. The Brier score decomposes to

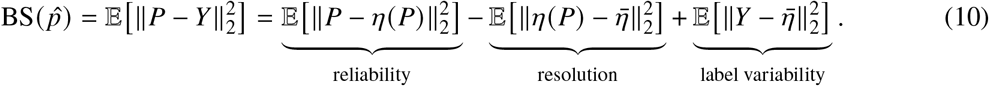

where *η*(*q*) = 𝔼 [*Y* | *P* = *q*] is the true class-frequency vector for instances receiving forecast *q* ∈ Δ^*K*−1^. Reliability measures how closely the predictive distribution matches empirical class frequencies. Resolution measures how strongly the forecast partitions the data into subsets with distinct class compositions relative to the global class prior 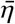. High resolution indicates that different forecast values correspond to meaningfully different conditional label distributions. The uncertainty term depends only on the class prior and is therefore fixed for a given dataset.

Evaluating calibration and resolution of the full multiclass distribution can be statistically challenging because it requires estimating conditional class frequencies over the entire simplex Δ^*K*−1^. A weaker and more practical notion therefore considers only the probability assigned to the predicted class [63]. Define the predicted class *G* = arg max_*j*_ *P* _*j*_, the confidence *C* = max_*j*_ *P* _*j*_ = *P*_*G*_, and the correctness indicator *A* = 𝟙 [*Y* = *G*], where 𝟙 [·] denotes the indicator function. This induces a binary probabilistic forecasting problem with forecast *C* ∈ [0, 1] and binary outcome *A* ∈ {0, 1}. Let *μ*(*c*) = *P*(*A* = 1 | *C* = *c*) be the outcome, ā = *P*(*A* = 1) be the overall accuracy.

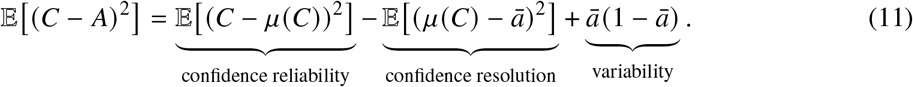

Here, confidence reliability measures whether confidences *C* are numerically calibrated, *P*(*Y* = *G* | *C* = *c*) = *P*(*A* = 1 | *C* = *c*) = *c*, while confidence resolution measures whether different confidences correspond to meaningfully different accuracies. High resolution indicates that predicted confidence is informative of outcomes, enabling separation between correct and incorrect predictions.

#### Frame-level Calibration Error and Resolution

Frame-level probabilities constitute the fundamental units of evidence that are accumulated during sequence decoding described in Sec. 2.6. Reliability and resolution can thus be assessed by examining the relationship between binned predicted confidence 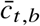, associated empirical accuracy 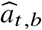, and mean accuracy 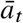 at this level.

We define confidence *c*_*t*_ for each frame and bin it into *B* equal-frequency bins with mean 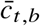.

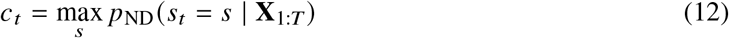

Given explicit frame-level annotations are unavailable for brain-to-text BCI, forced alignment provides a reference alignment for the ground-truth sentence **y**_1:*N*_. We define frame-level correctness as agreement between the prediction 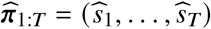 (Eq. 2), and reference 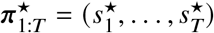 (Eq. 4) and calculate the empirical accuracy 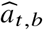 for each confidence bin ℬ_*b*_.

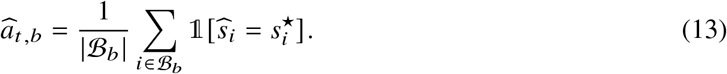

Expected calibration error (ECE) empirically quantifies the reliability of the predictive distribution by measuring the deviation between confidence and accuracy across confidence bins, approximating the calibration error 𝔼 (*C* − *μ*(*C*))^2^ in Eq. 11. Lower ECE indicates better calibration.

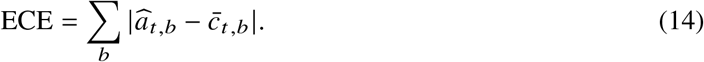

Resolution (RES) measures the dispersion of empirical accuracy across confidence bins relative to the mean accuracy 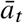, reflecting the resolution 𝔼 (*μ*(*C*) − ā)^2^ in Eq. 11. Higher RES indicates that uncertainty better informs error.

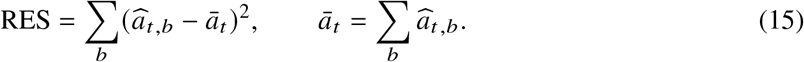

We use the area under the precision–recall curve (AUPR) to further quantify how well confidence discriminates between correct and incorrect predictions at frame or word levels, where higher AUPR ∈ [0, 1] indicates better separability.

#### Word-level Calibration Error and Resolution

Frame-level probabilities reflect the instantaneous uncertainty of neural decoding. Aggregating these probabilities over the duration of a word allows us to evaluate how uncertainty propagates from token-level predictions to semantically interpretable words. This analysis also helps diagnose brain-to-text BCI performance within the co-control framework, as language models operate primarily at the word level and word-level uncertainty is central to generating hypotheses 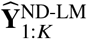 (Eq. 9; Sec. 2.6).

Each word corresponds to a contiguous span of tokens in time, we therefore define word-level correctness as agreement between the ground-truth *w* in sentence and the predicted word 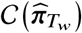 over the corresponding time span *T*_*w*_. Word-level confidence is obtained by aggregating frame-level *p*_ND_(*t*) over the same interval, yielding the confidence *c*_*w*_, accuracy 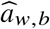, and mean confidence ā_*w*_.

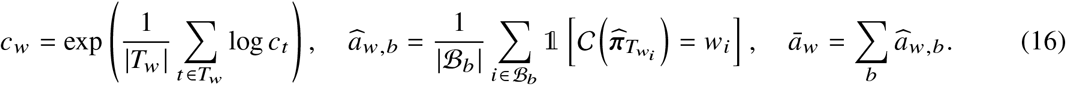

Using these quantities, ECE and RES are computed analogously to the frame-level:

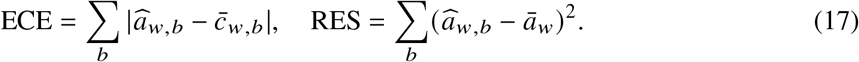

This hierarchical evaluation enables analysis of how uncertainty propagates from lower-level neural decoder predictions to structured outputs of interest, such as semantically meaningful words.

### 2.8 Oracle Simulations: Controlled Perturbations of Predictive Distribution

To isolate the role of probabilistic reliability (ECE, Eq. 14) and resolution (RES, Eq. 15) in *p*_ND_ from recognition accuracy, we introduce oracle-gated calibration to construct uncertainty-aware *p*_UA_ and overconfident *p*_OC_ model-output *p*_MO_ (Fig. 4A). These oracle simulations modify predictive distributions while preserving predicted tokens and the ranking of alternatives at each frame, enabling evaluation of how uncertainty quality alone affects decoding without altering accuracy (Fig. 2).

Oracle-gated calibration applies temperature scaling to *p*_MO_, redistributing probability mass with the temperature *τ*(·) conditioned on frame-level correctness 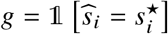 based on forced alignment.

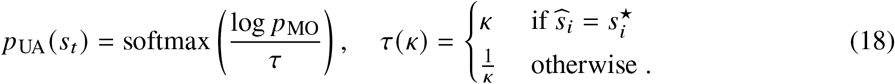

The gated temperature *τ*(*κ*) sharpens predictions for correct frames and flattens them for incorrect ones, improving calibration (alignment between confidence and correctness; ECE, Eq. 14) and resolution (discrimination between correct and incorrect predictions; RES, Eq. 15) at the same time. As a result, oracle-gated calibration provides an upper bound achievable via post-hoc calibration.

We also introduce an over-confident intervention, in which the predictive distribution is replaced by a degenerate distribution with Kronecker delta *δ*(·) on predicted token 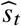 (Fig. 4A)

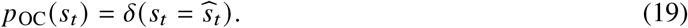

This transformation collapses the multinomial distribution to a vertex of the probability simplex Δ^*K*−1^, eliminating all uncertainty and reducing the decoder to deterministic predictions, worsening both ECE and RES. Consequently, this intervention represents a lower bound.

Effective decoding depends not only on the confidence of the most likely output, but also on the distribution of probability mass over alternative hypotheses. To isolate the role of this structure, we introduce a flat intervention that redistributes probability mass across competing hypotheses while preserving the predicted label. This transformation is applied to *p*_MO_ → *p*_MO−Flat_ and *p*_UA_ → *p*_UA−Flat_. The over-confident distribution *p*_OC_ is a special case for which *p*_OC_ = *p*_OC−Flat_.

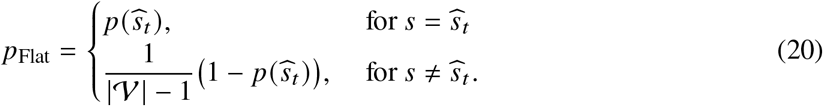

Oracle-gated calibration assumes access to framewise labels, which are obtained via forced alignment conditioned on the ground-truth transcript using the procedure described in Sec. 2.7. This oracle information is used exclusively for analysis and is not available during standard inference. Accordingly, oracle-gated calibration is not a realizable algorithm, but a diagnostic intervention used to assess how unreliable confidence limits decoding behavior under composition.

## 3 Results

In this section, we analyze the role of uncertainty in neural decoding at three levels. First, we evaluate the quality of the predictive distribution *p*_ND_ produced by neural decoders, showing that commonly used models exhibit systematic overconfidence (Sec. 3.1). Second, we investigate the functional role of uncertainty in sequence decoding, demonstrating how the structure of the predictive distribution causally influences hypothesis generation and integration during inference (Sec. 3.2). Finally, we examine the mechanisms that shape uncertainty estimation, showing that training objectives strongly influence the probabilistic semantics of neural decoder outputs (Sec. 3.2).

### 3.1 Predictive Distribution (*p*_ND_) from Neural Decoders Are Neither Reliable nor Informative

For neural decoders to convey useful uncertainty when decoding user intent, the predictive distribution (Eq. 1, Fig. 3A) must highlight situations in which neural evidence is weak, ambiguous, or unfamiliar resulting in predictions that are prone to error. In brain-to-text BCIs, such uncertainty signals guide interactions with other system components by encouraging diverse hypothesis generation, modulating reliance on language-model correction, or triggering user confirmation before executing commands in downstream assistive modules (Fig. 1). Consequently, evaluating the predictive distribution is necessary in addition to error metrics (e.g., PER, WER) on the decoded sentence.

**Figure 3:**
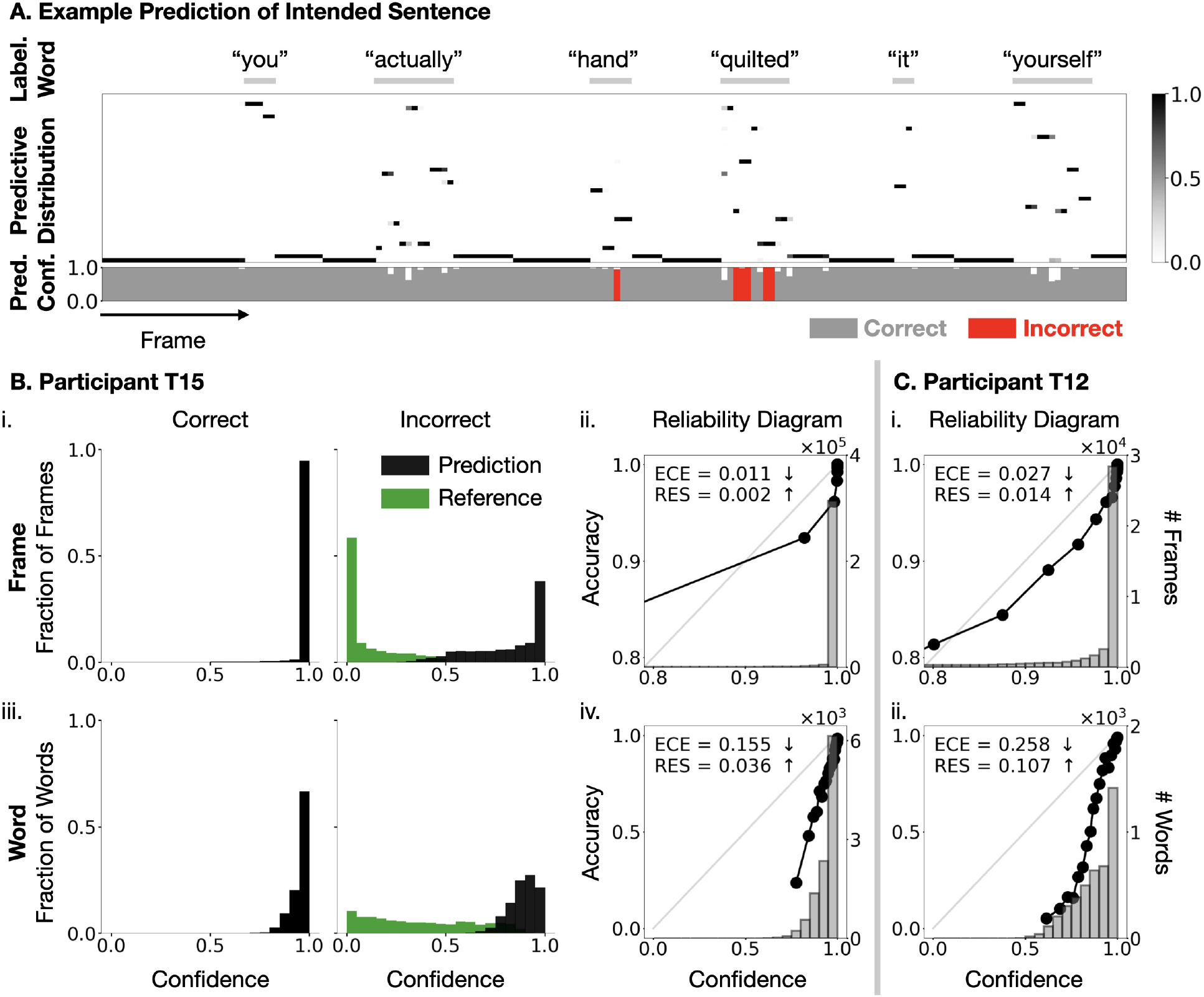
Neural decoder predictive distributions *p*_ND_ are over-confident. A) Example frame-level predictive distribution *p*_ND_ from a CTC-trained neural decoder for the sentence “you actually hand quilted it yourself.” The derived confidence *c*_*t*_ is shown below each frame; incorrect frames are highlighted in red. (B) Quantification of frame-level (i–ii) and word-level (iii–iv) uncertainty for participant T15. (i, iii) show confidence distributions for correct vs. incorrect predictions at the frame and word levels, respectively; green bars indicate the ground-truth class, revealing large margins in incorrect predictions and limited resolution. (ii, iv) show reliability diagrams, where binned confidence is compared to empirical accuracy (calibration curve, black dotted line), with the diagonal (grey line) indicating perfect calibration. At both levels, calibration curves lie below the diagonal, indicating systematic over-confidence (high ECE) and low resolution (low RES). (C) Reliability diagrams for participant T12 at the frame (i) and word (ii) levels, showing similar over-confidence.

The quality of probabilistic predictions is determined by two complementary properties: reliability and informativeness. Reliability (calibration) requires predicted confidence to match empirical correctness. For example, predictions assigned confidence 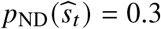 should be correct approximately 30% of the time, as measured by the expected calibration error ECE (Eq. 14). However, calibration alone is insufficient: a model with overall accuracy ā_*t*_ = 0.8 could trivially assign confidence *c*_*t*_ = 0.8 to every frame and achieve perfect calibration (ECE = 0) while providing no information about when errors are likely to occur. Informativeness is captured by resolution RES (Eq. 15), which measures variation in empirical correctness across confidence levels. Together, ECE and RES characterize probabilistic prediction quality under proper scoring rule decomposition (Eq. 10, 11) [63, 65, 79–82]. An effective neural decoder should therefore exhibit both low calibration error and high resolution, allowing the user or downstream modules to respond to localized uncertainty in decoded output.

Across all frames, neural decoders assign consistently high confidence (i.e., maximum predicted probability) to their predictions (Fig. 3B for T12, Fig. 3C for T15). The distribution of framewise confidence *c*_*t*_ (Eq. 12) is strongly concentrated near 1 for CTC-trained decoders (Sec. 2.3) on both participants. For participant T15, phoneme decoding achieves PER = 0.10. Considering only frames corresponding to phoneme predictions (excluding the 66% of frames predicting *ϵ* that primarily function as padding), 4.6% of frames are incorrect. However, these incorrect frames still receive high confidence (avg. 0.817 ± 0.188), only modestly lower than correct frames (avg. 0.970 ± 0.088), indicating limited separation between correct and incorrect outcomes (AUPR = 0.211 for T15 and 0.387 for T12). On error frames, the probability assigned to the ground-truth phoneme is much lower (avg. 0.096 ± 0.131), indicating that probability mass is concentrated on the predicted phoneme despite being incorrect, while suppressing plausible alternatives. Thus, although frame-level errors are rare, when they occur the model remains highly confident in incorrect predictions.

To evaluate whether *p*_ND_ are probabilistically meaningful, we analyze calibration using reliability diagrams and quantify ECE (Eq. 14) and RES (Eq. 15). Reliability diagrams reveal systematic overconfidence: predictions assigned high confidence frequently exhibit lower empirical accuracy than expected under ideal calibration (Fig. 3B(ii), Fig. 3C(i)). Quantitatively, the neural decoder exhibits measurable calibration error (T12: ECE = 0.027, T15: ECE = 0.011), while resolution remains low (T12: RES = 0.014, T15: RES = 0.002), indicating limited variation in empirical correctness across confidence levels. These metrics depend on label distribution (Sec. 2.7) and are not directly comparable across datasets. Together, these results indicate that neural decoder prediction *p*_ND_ provides unreliable and uninformative uncertainty estimates at the frame level.

This overconfidence propagates when probabilities are aggregated over time (Fig. 3B(iii)-(iv), Fig. 3C(ii)). At the word level, incorrect predictions remain associated with high confidence (T12: 0.930 ± 0.071 for correct vs. 0.782 ± 0.099 for incorrect; T15: 0.958 ± 0.047 for correct vs. 0.889 ± 0.072 for incorrect), yielding weak separation between correct and incorrect word predictions (AUPR = 0.829 for T12 and 0.547 for T15). Reliability and informativeness also deteriorate with high calibration error (ECE = 0.258 for T12; 0.155 for T15) and low resolution (RES = 0.107 for T12; 0.036 for T15). These results indicate that frame-level overconfidence carries over to word-level uncertainty.

Together, these analyses reveal that neural decoder predictive distributions *p*_ND_ are neither reliable nor informative. At the frame level, confidence fails to meaningfully distinguish correct from incorrect phoneme predictions, with probability mass concentrated on the predicted phoneme and little support for alternatives (Fig. 3 B(ii), Fig. 3 C(i)). When aggregated over time, this overconfidence propagates and obscures the distinction between correct and erroneous words (Fig. 3 B(iv), Fig. 3 C(ii)). These results expose two probabilistic failure modes in current neural decoders: frame-level overconfidence and temporal propagation of overconfidence during sequence decoding. As a result, predictive distribution provide little information about errors and fail to reliably convey uncertainty to downstream components or the user. In the next section, we examine how predictive distributions influence decoding behavior, revealing their role in probabilistic co-control brain-to-text BCIs.

### 3.2 Oracle Simulations Reveal the Causal Role of Predictive Distribution in Decoding

Brain-to-text decoding operates entirely on predictive distribution *p*_ND_ (Eq. 1), which downstream modules interpret as uncertainty-aware control signals (Sec. 2.6). Results in Sec. 3.1 show that existing neural decoders fail to express reliable and informative uncertainty: confidence remains high even when predictions are incorrect. We therefore ask a more fundamental question: to what extent does the predictive distribution alone constrain decoding performance, independent of accuracy?

To isolate this effect, starting from the model predictive distribution *p*_MO_ analyzed in Sec. 3.1, we generate two interventions, an over-confident distribution *p*_OC_ and an uncertainty-aware distribution *p*_UA_, while holding all other factors fixed (detailed in Sec. 2.8; Fig. 4A). First, we collapse the distribution so that all the probability mass is assigned to the predicted label at every frame, producing the over-confident distribution *p*_OC_ (Eq. 19). Because this representation contains no usable uncertainty information, it serves as a lower bound on the utility of probabilistic prediction. Second, we simulate an uncertainty-aware distribution *p*_UA_ (Eq. 18) using oracle-gated temperature scaling. This intervention increases the margin between the confidence assigned to correct and incorrect frames while preserving the ranking of alternative tokens. In this way, *p*_UA_ approximates an upper bound on useful uncertainty estimation. The three distributions *p*_MO_, *p*_OC_, and *p*_UA_ therefore represent alternative realizations of the neural decoder prediction *p*_ND_. We propagate these probabilities through the multi-stage decoding pipeline and examine how they influence hypothesis generation 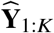 (Eq. 9) and language-model integration 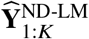 (Eq. 9).

We first examine how different predictive distribution constructions affect the statistical properties of frame-level confidence. Under the uncertainty-aware intervention *p*_UA_, predicted confidence exhibits a bimodal distribution that separates correct from incorrect predictions. Reliability diagrams (Fig. 4B.(i)-(iii)) show that UA closely tracks ideal calibration, whereas the model output MO and over-confident OC baselines systematically overestimate accuracy to varying degrees. This separation between correct and incorrect predictions separation substantially increases resolution (RES, Eq. 15), indicating improved discriminability between correct and incorrect predictions. Resolution values are reported in the order ⟨*U A, MO, OC*⟩; for T12: ⟨0.066, 0.014, 0.000⟩, and for T15: ⟨0.005, 0.002, 0.000⟩. At the same time, UA improves the correspondence between predicted confidence and empirical accuracy, reducing calibration error (ECE, Eq. 14) (T12: ⟨0.014, 0.027, 0.084⟩; T15: ⟨0.000, 0.011, 0.016⟩). Because task complexity is fixed within each dataset, RES and ECE are directly comparable across the *OC, MO*, and *U A*.

Importantly, these improvements persist across temporal aggregation. For word-level predicted confidence (Sec. 2.7), UA again yields higher RES and lower ECE than *MO* and *OC* (Fig. 4B(iv)-(vi)). Consequently, word-level uncertainty becomes a meaningful indicator of decoding errors: words assigned low confidence under UA are highly likely to be incorrect, achieving strong error identification performance (AUPR = 0.992 for T12, 0.962 for T15). This demonstrates that improved probability quality translates directly into usable uncertainty signals at the word level.

Reliable probabilities also produce immediate effects on the search dynamics of decoding. During hypothesis generation, beam search driven by UA explores a broader and more structured hypothesis space. Although frame-level recognition accuracy is unchanged, the correct sequence appears more frequently among lower-ranked hypotheses, with gains in top-*K* Δ_OC-UA_: 0.008 (*K* = 1), 0.024 (*K* = 5), and 0.040 (*K* = 100) for T15; and 0.014 (*K* = 1), 0.030 (*K* = 5), and 0.049 (*K* = 100) for T12 (Fig. 4B(vii), Fig. 4C(i)). These improvements indicate that informative uncertainty preserves competing hypotheses during search rather than prematurely collapsing onto a single hypothesis.

The benefits extend further during hypothesis integration with the language model. Because UA maintains greater hypothesis diversity, language-model re-ranking operates more effectively and more consistently promotes the ground-truth sequence. This yields systematic WER reductions across candidate sets (Δ_OC-UA_ = ⟨0.025, 0.012, 0.003⟩ for T15; ⟨0.079, 0.066, 0.042⟩ for T12; Fig. 4B(vii), Fig. 4C(ii)). Not all gains from hypothesis generation propagate to integration, as lexicon constraints prune invalid phoneme sequences and alter the effective predictive distribution by removing some high-confidence but incorrect hypotheses. Nevertheless, the remaining improvements demonstrate that reliable uncertainty enables more effective interaction between neural decoding and language-model correction.

Together, these results reveal a third failure mode in current neural decoding systems: unreliable probability estimates can simultaneously impair both hypothesis generation and hypothesis integration in decoding (Fig. 2, 4B(vii)-(viii), Fig. 4C(i)-(ii)). When uncertainty is poorly represented, beam search collapses prematurely, limiting alternative hypotheses, while language-model integration receives unreliable signals for re-ranking (Fig. 2). Comparisons across simulated outputs (UA, MO, OC) show that improvements in recognition accuracy (PER) do not necessarily translate to end-to-end gains (WER). Our analysis shows that WER of system output depends jointly on recognition accuracy and the quality of uncertainty estimates, such that improvements in PER can be offset by poorly calibrated or uninformative uncertainty. Reliable and structured uncertainty is therefore not merely desirable, but functionally necessary for effective interaction between neural decoding and behavior modules in probabilistic co-control brain-to-text systems. This oracle analysis highlights a substantial performance gap that can be addressed through improved uncertainty estimation, motivating the novel neural decoder design introduced in the following sections.

### 3.3 Balancing Alignment Learning and Classification Produces Calibrated Probabilities

Neural decoders in brain-to-text BCIs are trained to optimize phoneme recognition under weak supervision, yet their predictive distribution are subsequently used as uncertainty signals by downstream modules. This raises a central question: can predictive distribution be learned that faithfully reflect uncertainty while preserving decoding accuracy? A fundamental challenge is that neither framewise labels nor uncertainty are directly observable, as the alignment between neural activity and phoneme is latent. The CTC loss marginalizes over all valid alignments (Sec. 2.3), often producing “peaky alignments” that concentrate frame-level probability mass on a single predicted token. This implicitly selects a dominant alignment during learning (Sec. 2.4), reinforcing iterative fitting to that alignment hypothesis and thereby introducing artificial certainty [74, 75]. In this setting, the over-confidence required to stabilize alignment may suppress the proper expression of frame-wise uncertainty arising from model uncertainty, ambiguous neural evidence, or label noise.

If over-confidence arises primarily from resolving alignment ambiguity, then stabilizing the alignment target should improve the informativeness and reliability of predictive distribution. Conversely, if uncertainty is dominated by irreducible variability in neural evidence, stabilizing the alignment target would not improve, and may degrade decoding. To test this hypothesis, we introduce a two-stage training strategy in which a CTC model infers pseudo-alignments 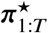 (Eq. 4), which are then fixed to train a CE model (Sec. 2.4), explicitly decoupling alignment inference from classification. The resulting models, *p*_CE_ and *p*_CTC_, provide a controlled contrast between alignment-marginalized learning and fixed-target classification under otherwise identical conditions (Fig. 5A,B).

**Figure 4:**
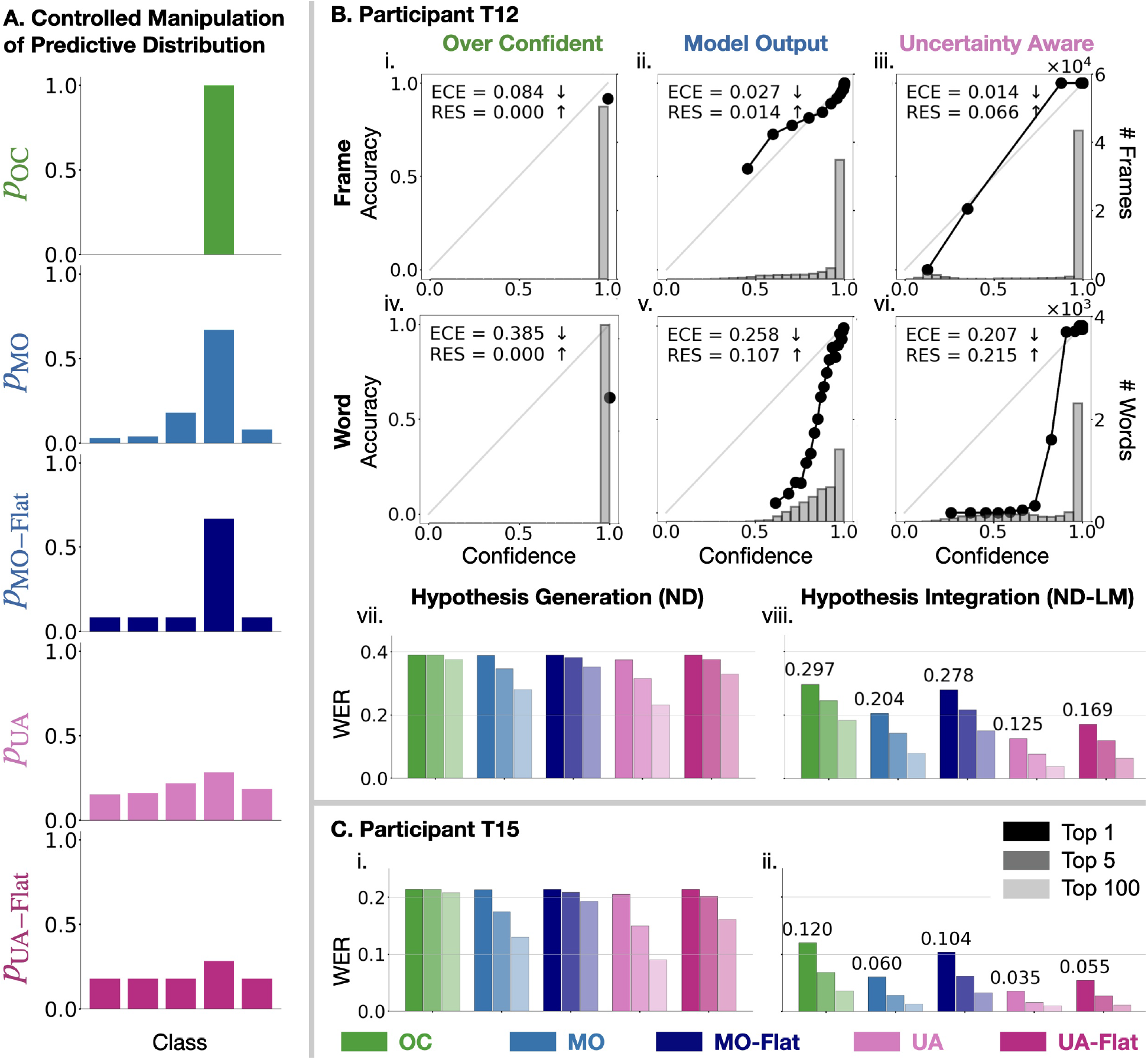
Oracle interventions reveal the causal effect of predictive distributions on decoding, independent of accuracy. (A) Illustration of controlled manipulations of the predictive distribution *p*_ND_ (restricted to five classes for visualization), showing over-confident (*p*_OC_), uncertainty-aware (*p*_UA_), and flattened variants (*p*_MO−Flat_, *p*_UA−Flat_), compared to the original model output (*p*_MO_). (B) Comparison of oracle interventions (*p*_OC_, *p*_UA_) and flattened variants (*p*_MO−Flat_, *p*_UA−Flat_) to the original neural decoder prediction (*p*_MO_) for participant T12. Reliability diagrams (introduced in Fig. 3) and summary metrics (ECE, RES) show substantial differences in how confidence reflects accuracy across *p*_OC_, *p*_MO_, *p*_UA_ at both the frame (i–iii) and word (iv–vi) levels. All manipulations preserve identical frame-wise errors, yet yield different decoding performance for (vii) sequence decoding based on *p*_ND_ alone and (viii) hypothesis integration with the language model. Shaded regions indicate WER for top-1, top-2, and top-100 hypotheses. (C) Decoding performance under the same manipulations applied to *p*_MO_ for participant T15, showing similar dependence of performance on the predictive distribution. Format follows (B).

**Figure 5:**
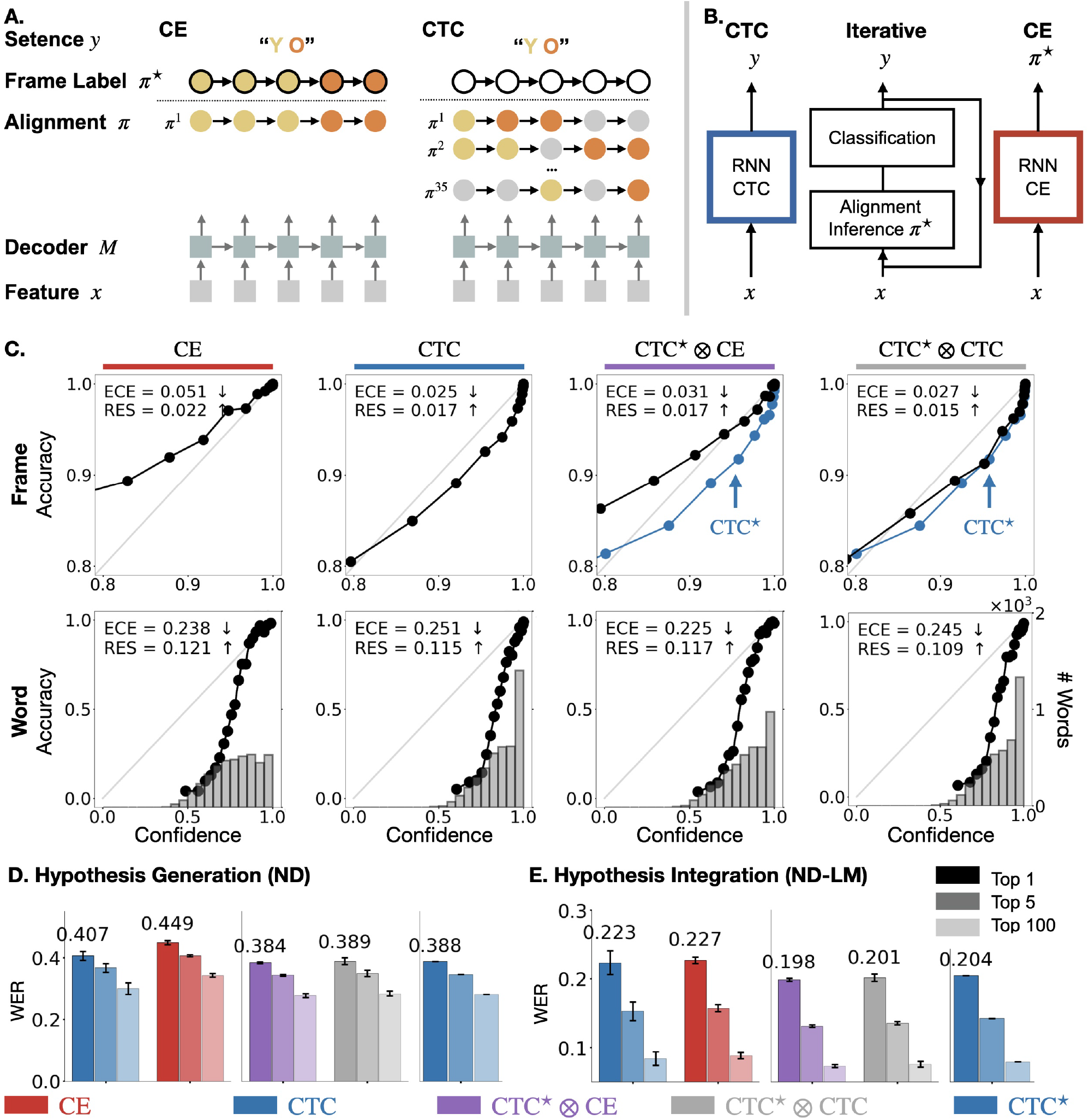
Fusing CTC and CE predictive distributions (CTC★ ⊗ CE) yields more calibrated predictions. Conceptual comparison of CE and CTC training. (A) Valid alignment hypotheses under CTC and CE for the example sentence “YO”. (B) Training schemes for CTC, which marginalizes over all valid alignments for the target sequence **y**, and CE, which is trained with respect to a fixed alignment ***π***^★^. (C) Comparison of frame- and word-level reliability diagrams (introduced in Fig. 3) and summary metrics (ECE, RES) for *p*_CE_ and *p*_CTC_ across random initialization, and fused *p*CTC★ ⊗ CE and *p*CTC★ ⊗ CTC using the best CTC model CTC★. Decoding performance across models. (D) Hypothesis generation and (E) language model integration, showing improved top-1 and top-5 performance with CTC★ ⊗ CE.

We first contrast the probabilistic behavior of neural decoders trained with CTC and CE objectives. Using identical architectures, training data, and alignment hypotheses, we isolate the effect of the training objective on predictive distribution (Sec. 2.3; 2.4). CTC-trained models achieve higher frame-level accuracy by a small margin (CTC:0.848 ± 0.005 vs CE:0.832 ± 0.002), leading to slightly lower PER (CTC:0.212±0.009 vs CE:0.232±0.002). Despite this moderate advantage in recognition accuracy, CTC models produce systematically over-confident probabilities (Sec. 3.1). In contrast, CE-trained models exhibit markedly different probability behavior (Fig. 5C). In particular, CE achieves higher resolution (RES: CTC 0.017 vs CE 0.022), indicating improved discrimination between correct and incorrect predictions (AUPR: CTC 0.426 ± 0.016 vs CE 0.469 ± 0.006). Although CE exhibits higher calibration error (ECE: CTC 0.025 vs CE 0.051), its reliability curve lies above the diagonal, revealing systematic under-confidence rather than over-confidence. Correspondingly, the confidence of *p*_CE_ spans a wider dynamic range and more closely tracks empirical correctness. These results demonstrate that training objectives can induce qualitatively different uncertainty behaviors beyond simple over-confidence, even while maintaining comparable recognition accuracy.

Over-confident predictive distribution limit its functional use during decoding. In particular, collapsed beam degrade hypothesis generation by suppressing lower-ranked candidates, while extreme token scores hinder effective hypothesis integration with the language model (Sec. 3.2; Fig. 2). We therefore ask whether the observed differences in probabilistic behavior between CTC and CE models impact downstream decoding. Although improved uncertainty estimation does not substantially change the quality of lower-ranked hypotheses (top-100 WER improves top-1 by Δ_WER_ = 0.108 for CE vs 0.108 for CTC), it produces more informative hypothesis scores. As a result, beam search benefits more from language model integration when using *p*_CE_, yielding larger WER reductions from pre-(Fig. 5D) to post-integration (Fig. 5E) (ΔWER: 0.222 for *p*_CE_ vs 0.184 for *p*_CTC_). These results show that uncertainty estimation has functional consequences for decoding by shaping the hypothesis set and their scores, which determine the effectiveness of language-model re-ranking.

Motivated by the complementary behaviors of CTC and CE models, we evaluate a controlled fusion using the best-performing CTC model CTC^★^ (selected based on validation performance) and compare CTC^★^ ⊗ CE, CTC^★^ ⊗ CE against CTC^★^ alone (Sec. 2.5). A single interpolation parameter *γ* continuously trades off between alignment-dominant (*p*CTC★) and explicit uncertainty (*p*_CE_) prediction. As a control, we also fuse *p*CTC★ and *p*_CTC_. Across all *γ*, fused models reduce frame- and word-level calibration error, increase effective resolution, and improve decoding utility relative to either model alone. Performance peaks at an intermediate *γ* = 0.5 (ΔCTC★+CE,CTC★ = 0.006, 0.011, 0.006 for *K* = 1, 5, 100) (Fig. 5C-E). These improvements indicate that fusing model predictions allows disagreement between models to act as a more reliable and informative probability estimate. Combining models with distinct inductive biases may further amplify this benefit.

Together, these results show that training objectives shape the semantics of predictive distribution. Although CTC-trained models achieve slightly higher frame-level accuracy, they produce systematically over-confident predictions consistent with alignment–classification coupling in CTC. By stabilizing the alignment target, CE training decouples alignment from prediction, yielding predictive distribution whose uncertainty better tracks empirical correctness. Model fusion further leverages disagreement between models, especially those with distinct inductive biases such as CTC and CE to improve uncertainty estimates. Taken together, these findings suggest that uncertainty behavior in neural decoders is not determined solely by model architecture or data statistics, but can be systematically shaped by the choice of training objective.

## 4 Discussion

BCI control signals typically contain errors because neural input features are inherently variable and rarely provide complete evidence of the user’s intended behavior. To support effective and safety-critical BCIs, neural decoders must therefore do more than produce a single best estimate of user intent: they must also communicate uncertainty about their predictions. Most prior work has focused on improving end-to-end accuracy metrics such as word error rate [11,12,35–39,57]. However, neural decoders operate as probabilistic inference modules within co-control BCI systems, where predictive distribution function as uncertainty signals that guide downstream components, including language-model integration and user feedback (Fig. 1).

Our results show that neural decoder predictive distribution (Sec. 2.2) cannot be assumed to faithfully represent uncertainty. Widely used neural decoders trained with the CTC objective [11, 12, 35–39, 57, 73] exhibit systematic overconfidence, assigning near-deterministic confidence even when predictions are incorrect (Sec. 3.1). This overconfidence suppresses alternative hypotheses during sequence decoding and distorts hypothesis arbitration with language models (Sec. 3.2; 3.3). In the following discussion sections, we synthesize these findings to characterize the consequences of these failure modes for probabilistic inference (Sec. 4.1), connect the mechanisms underlying overconfident predictions to concrete directions for improving uncertainty estimation (Sec. 4.2), and articulate broader implications for the design of effective co-control and its extension to other BCI tasks (Sec. 4.3). More broadly, our work highlights that relying solely on end-to-end decoding accuracy can obscure critical properties of the neural decoder’s probabilistic outputs, potentially biasing system design toward specific AI models or evaluation settings rather than producing reliable uncertainty signals that can support multiple downstream modules. By opening this “black box,” we aim to characterize and evaluate the implicit control signal, decoder predicted uncertainty, that governs interaction between neural decoding and AI components in next-generation BCI systems.

### 4.1 Neural Decoders Do Not Provide Informative and Reliable Estimates of Uncertainty

Analysis of predictive distribution from widely used CTC neural decoders for brain-to-text BCI [11, 12, 35–39,57] reveals a consistent probabilistic structure across subject (T12 [12] and T15 [35]). Specifically, neural decoders exhibit three coupled probabilistic failure modes: predictive distribution collapse toward near-deterministic frame-level predictions; this overconfidence propagates through time, aggregating into overconfident word-level predictions; and the resulting certainty suppresses alternative hypotheses during sequence decoding and distorts the language-model integration.

First, frame-level overconfidence causes the predictive distribution to concentrate on the predicted phoneme even when the prediction is prone to error. This near-deterministic behavior prevents uncertainty from being localized within a sequence, yielding poor reliability (high ECE) and low informativeness (low RES) as quantified by the proposed metrics (Sec. 2.7; Fig. 3). Consequently, downstream components cannot distinguish frames supported by strong neural evidence from those arising from ambiguous or noisy input, preventing adaptive responses to local uncertainty.

Second, this miscalibration propagates temporally through the decoding hierarchy. Overconfident frame predictions accumulate into overconfident word-level predictions (Sec. 2.7, 3.1; Fig. 3). Incorrect words therefore frequently appear both wrong and certain, making them difficult to flag for user confirmation or correction. This failure is particularly consequential for low-frequency yet semantically informative words, which are known to exhibit higher error rates in brain-to-text decoding [83]. When such errors are accompanied by high confidence, both users and downstream modules lose the ability to efficiently detect and correct mistakes.

Third, overconfident neural predictions can cause multi-stage decoding to fail by constraining both hypothesis generation and hypothesis integration (Fig. 2). When predictive distributions concentrate excessively on a single token, beam search prematurely suppresses alternative hypotheses, thus producing a narrow candidate set and limiting sequence diversity (Fig. 4 B,C), a failure mode widely observed in general sequence decoding systems [84–90]. This restricted hypothesis space reduces the effectiveness of language-model integration, which relies on graded score differences among plausible alternatives to perform contextual re-ranking and error correction (Fig. 4 B,C). Notably, sentences with higher phoneme error rates benefit most from calibrated uncertainty (Fig. 6 A). Although these cases contain weaker contextual information, localized uncertainty preserves candidate hypotheses that would otherwise be pruned during decoding, allowing the language model to recover from errors more effectively. This helps explain why improvements in phoneme error rate (PER) do not always translate to lower word error rate (WER) in integrated systems [57]. Scalar accuracy metrics capture correctness, but not how predictive uncertainty structures intermediate hypotheses and module interactions in multi-stage inference (Sec. 3.2).

**Figure 6:**
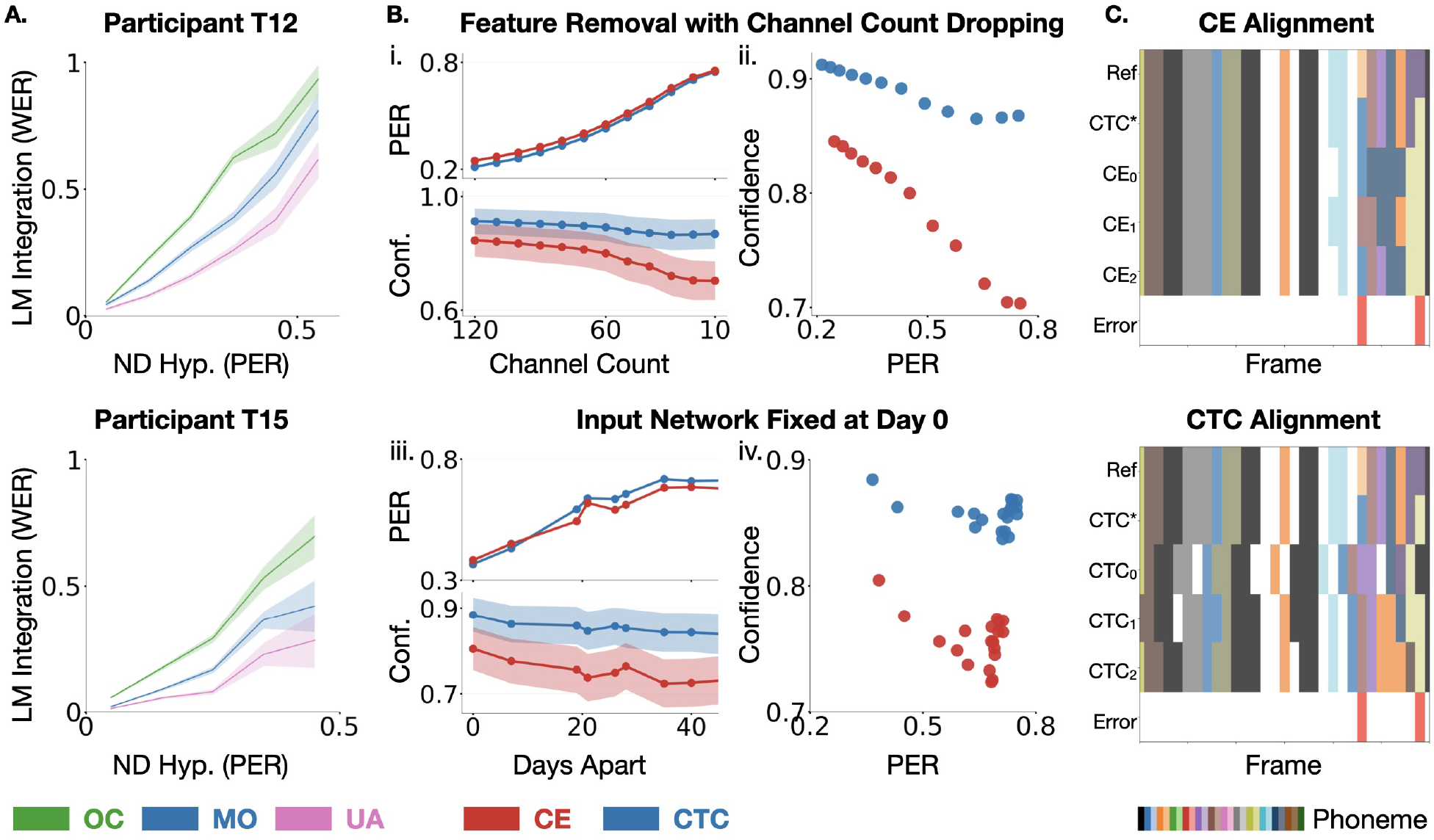
Predicted uncertainty should reflect evidence in the neural signal. (A) Effect of LM integration (WER) across sentences at different decoding error levels (PER), showing a persistent gap between oracle over-confident (*p*_OC_, green) and uncertainty-aware (*p*_UA_, pink) conditions relative to the model output (*p*_MO_, blue). (B) Decoding performance (PER) and uncertainty (confidence) as a function of information degradation: (i–ii) varying the number of electrodes used for offline decoding, and (iii–iv) increasing temporal drift from a model with input network trained on Day 0. In regimes with substantial PER degradation (i, iii), *p*_CTC_ remains over-confident while *p*_CE_ reduces confidence in proportion to performance loss. (C) Example alignments for a sentence under randomly initialized CTC and CE models, compared to a reference alignment ***π***^★^ (Ref). Errors (CTC^★^ vs Ref) illustrate that correct predictions can still yield inconsistent phoneme alignments, reflecting alignment uncertainty.

These findings suggest that neural decoder predictive distributions act as control signals governing decoding behavior. When these signals are miscalibrated, a system may appear accurate while silently failing during ambiguous predictions. Improving the reliability and informativeness of decoder-predicted uncertainty is therefore essential for enabling robust hypothesis generation, effective language-model integration, and ultimately safer co-control BCI systems.

### 4.2 Mechanisms of Miscalibrated Uncertainty and Design Implications

An ideal neural decoder should learn to express uncertainty that is localized in time and reflects the strength of neural evidence supporting ongoing behavior. Through controlled comparisons between neural decoders trained with CTC (Sec. 2.3) and CE (Sec. 2.4) objectives, we find that training objectives strongly influence the uncertainty representation in prediction. The CTC objective couples alignment and classification, encouraging overconfident predictions that resolve alignment ambiguity. In contrast, decoupling alignment via CE training preserves accuracy while improving probability resolution (Sec. 3.3), yielding more informative uncertainty estimates. Training objectives serve as a key design lever for shaping uncertainty estimation, in addition to determining recognition accuracy.

In brain–computer interfaces, single-trial neural signals are inherently noisy and variable [91–93], and user control signals fluctuate across attempts due to factors such as fatigue, feedback, and task familiarity [8, 94–103]. Consequently, reliable uncertainty estimates should adapt to variations in the quality of available neural evidence. However, natural variability is difficult to isolate without time-aligned ground-truth behavior. To address this challenge, we introduce controlled signal degradation experiments that simulate two common sources of performance decline: progressive channel dropout and cross-session non-stationarity arising from the removal of adaptive input updates. Under these conditions, CTC-trained decoders remain highly confident despite substantial loss of information, whereas CE-trained models exhibit increased uncertainty as accuracy deteriorates (Fig. 6B). These results suggest that alignment-driven objectives can override evidence, producing confidence collapse rather than evidence-aligned uncertainty. This distinction becomes particularly important in practical BCI scenarios where neural signals vary substantially across users and recording conditions, such as differences in clinical severity [12, 35, 37, 52] or different modes of control (e.g., inner speech [38] versus attempted speech [12, 35,37,52]). In such settings, reliable and evidence-based uncertainty estimation is critical for maintaining robust system behavior.

Beyond modifying training objectives, another approach to improving uncertainty estimation is to leverage model ensembles [70]. By combining independently trained models, ensembles capture uncertainty induced by distinct optimization trajectories. In our experiments, combining CTC and CE models yields the largest improvements in both uncertainty estimation and decoding accuracy. Analysis across random initializations reveals that CTC models frequently disagree in their inferred alignments while still predicting the same phoneme sequence, indicating inherent alignment uncertainty in the learned representations [104]. In contrast, CE models share a fixed alignment but often disagree on specific token predictions when segments are likely to contain errors. These complementary disagreement patterns help explain why combining models trained with different objectives improves uncertainty estimation. This perspective may also explain why several recent systems with CTC-based neural decoders perform hypothesis ensembling only after language-model integration and rely on additional large models for rescoring or rewriting [47,53,57,58]. While effective, this approach often incurs substantial computational cost and shifts increasing responsibility to large language models, which resolve errors primarily using semantic context rather than uncertainty conveyed by the neural evidence of user intent.

### 4.3 Calibrated Uncertainty As A Foundation For Co-control

The central limitation in next-generation co-control BCIs may not be decoding accuracy, but the faithful representation of uncertainty. While accuracy measures performance in static, monolithic integrations of neural decoders and behavioral modules, uncertainty estimation determines how effectively neural decoders collaborate with downstream modules under increasing complexity, partial observability, and evolving conditions. This shift in perspective reframes uncertainty as a system-level quantity that links multiple stages of BCI development, from the encoding and decoding of neural signals to the integration, validation, and execution of predictions by downstream modules.

Uncertainty can take different forms across modalities. In discrete decoding tasks such as brain-to-text, it is expressed as predictive distributions over states, whereas in continuous motor BCIs (e.g., arm [5, 6, 8–10, 14] and finger movement [105–107]), it naturally appears as variance over movement trajectories. Across these settings, uncertainty is not merely descriptive, rather it functions as a control signal through which neural evidence is communicated, interpreted, and integrated with downstream modules. However, reliable uncertainty does not arise automatically from standard training paradigms. Although most objectives produce predictive distributions or variance, the reliability of these uncertainty estimates is shaped by the objective, often leading to misalignment with the underlying neural evidence. This suggests that uncertainty must be explicitly modeled and evaluated. A key direction is to develop uncertainty-aware objectives that align predictive uncertainty with the structure of neural variability, for example by enforcing consistency between uncertainty estimates and behavior-relevant neural latent dynamics. Such alignment may improve reliability at fine temporal scales and enable more robust system behavior. As uncertainty-aware decoders mature, uncertainty signals may increasingly guide decision-making under constraints, including risk-sensitive actions, user–BCI co-adaptation [95, 108–111], and interaction with multiple modules. Uncertainty not only modulates control, but may define new modes of interaction in translational BCIs.

Taken together, these directions reinforce the role of uncertainty as a central interface for coordinating interaction across modules in co-control systems. Viewed through this lens, the neural decoder becomes a collaborative partner rather than a rigid translator of neural activity, enabling co-control systems that protect users from errors under weak or ambiguous neural signals. By prioritizing calibrated trust and principled deferral, such systems can function as reliable, life-enhancing extensions of a person’s communicative intent while maintaining robustness in safety-critical settings. More broadly, this perspective suggests a development paradigm for BCIs in which uncertainty serves as the connective layer linking neurophysiological variability, representation and decoder design, robustness evaluation, and system-level integration. In this view, uncertainty is not merely a property of prediction, but the mechanism, and ultimately the interface, through which neural intent is translated, evaluated, and integrated with intelligent systems, enabling safe, adaptive, and interpretable co-control across increasingly complex behaviors.

## Notes

### Competing Interest Statement

The authors declare the following financial interests which may be considered as potential competing interests with the work reported in this paper: VG holds shares in Neuralink Corp. and is Chief Scientific Officer and an options holder at Paradromics, Inc.

### Funding

- National Institutes of Health: R01DC018446
- U.S. National Science Foundation: 2223822

## References

[1] Krishna V. Shenoy and Byron M. Yu. Brain–machine interfaces. In Eric R. Kandel, John D. Koester, Sarah H. Mack, and Steven A. Siegelbaum, editors, Principles of Neural Science, chapter 39. McGraw-Hill, 6 edition, 2021.

[2] Jacob T Robinson, Sumner L Norman, Matthew R Angle, Timothy G Constandinou, Timothy Denison, John P Donoghue, Ryan M Field, Andreas Forsland, Sid Kouider, Josédel R Millán, et al. An application-based taxonomy for brain–computer interfaces. Nature Biomedical Engineering, 9(6):789–791, 2025.

[3] Mariska J Vansteensel, Eran Klein, Ghislaine van Thiel, Michael Gaytant, Zachary Simmons, Jonathan R Wolpaw, and Theresa M Vaughan. Towards clinical application of implantable brain– computer interfaces for people with late-stage als: medical and ethical considerations. Journal of Neurology, 270(3):1323–1336, 2023.

[4] Gerwin Schalk, Peter Brunner, Brendan Z Allison, Surjo R Soekadar, Cuntai Guan, Tim Denison, Jörn Rickert, and Kai J Miller. Translation of neurotechnologies. Nature Reviews Bioengineering, 2(8):637–652, 2024.

[5] Frank H. Guenther, Jonathan S. Brumberg, E. Joseph Wright, Alfonso Nieto-Castanon, Jason A. Tourville, Mikhail Panko, Robert Law, Steven A. Siebert, Jess L. Bartels, Dinal S. Andreasen, Princewill Ehirim, Hui Mao, and Philip R. Kennedy. A wireless brain-machine interface for real-time speech synthesis. PloS one, 4(12):e8218, 2009.

[6] Daniel Bacher, Beata Jarosiewicz, Nicolas Y Masse, Sergey D Stavisky, John D Simeral, Katherine Newell, Erin M Oakley, Sydney S Cash, Gerhard Friehs, and Leigh R Hochberg. Neural point-and-click communication by a person with incomplete locked-in syndrome. Neurorehabilitation and neural repair, 29(5):462–471, 2015.

[7] Beata Jarosiewicz, Anish A. Sarma, Daniel Bacher, Nicolas Y. Masse, John D. Simeral, Brittany Sorice, Erin M. Oakley, Christine Blabe, Chethan Pandarinath, Vikash Gilja, Sydney S. Cash, Emad N. Eskandar, Gerhard Friehs, Jaimie M. Henderson, Krishna V. Shenoy, John P. Donoghue, and Leigh R. Hochberg. Virtual typing by people with tetraplegia using a self-calibrating intracortical brain-computer interface. Science Translational Medicine, 7(313):313ra179, 2015.

[8] Vikash Gilja, Paul Nuyujukian, Cindy A Chestek, John P Cunningham, Byron M Yu, Joline M Fan, Mark M Churchland, Matthew T Kaufman, Jonathan C Kao, Stephen I Ryu, and Krishna V Shenoy. A high-performance neural prosthesis enabled by control algorithm design. Nature Neuroscience, 15(12):1752–1757, 2012.

[9] Vikash Gilja, Chethan Pandarinath, Christine H Blabe, Paul Nuyujukian, John D Simeral, Anish A Sarma, Brittany L Sorice, János A Perge, Beata Jarosiewicz, Leigh R Hochberg, Krishna V Shenoy, and Jaimie M Henderson. Clinical translation of a high-performance neural prosthesis. Nature Medicine, 21(10):1142–1145, 2015.

[10] Chethan Pandarinath, Paul Nuyujukian, Christine H Blabe, Brittany L Sorice, Jad Saab, Francis R Willett, Leigh R Hochberg, Krishna V Shenoy, and Jaimie M Henderson. High performance communication by people with paralysis using an intracortical brain-computer interface. elife, 6:e18554, 2017.

[11] Francis R Willett, Donald T Avansino, Leigh R Hochberg, Jaimie M Henderson, and Krishna V Shenoy. High-performance brain-to-text communication via handwriting. Nature, 593(7858):249– 254, 2021.

[12] Francis R Willett, Erin M Kunz, Chaofei Fan, Donald T Avansino, Guy H Wilson, Eun Young Choi, Foram Kamdar, Matthew F Glasser, Leigh R Hochberg, Shaul Druckmann, Krishna V. Shenoy, and Jaimie M. Henderson. A high-performance speech neuroprosthesis. Nature, 620(7976):1031– 1036, 2023.

[13] Philip R Kennedy, Roy AE Bakay, Melody M Moore, Kim Adams, and John Goldwaithe. Direct control of a computer from the human central nervous system. IEEE Transactions on rehabilitation engineering, 8(2):198–202, 2000.

[14] Paul Nuyujukian, Jose Albites Sanabria, Jad Saab, Chethan Pandarinath, Beata Jarosiewicz, Christine H Blabe, Brian Franco, Stephen T Mernoff, Emad N Eskandar, John D Simeral, Leigh R. Hochberg, Krishna V. Shenoy, and Jaimie M. Henderson. Cortical control of a tablet computer by people with paralysis. PloS one, 13(11):e0204566, 2018.

[15] Leigh R Hochberg, Mijail D Serruya, Gerhard M Friehs, Jon A Mukand, Maryam Saleh, Abraham H Caplan, Almut Branner, David Chen, Richard D Penn, and John P Donoghue. Neuronal ensemble control of prosthetic devices by a human with tetraplegia. Nature, 442(7099):164–171, 2006.

[16] Leigh R Hochberg, Daniel Bacher, Beata Jarosiewicz, Nicolas Y Masse, John D Simeral, Joern Vogel, Sami Haddadin, Jie Liu, Sydney S Cash, Patrick Van Der Smagt, and John P. Donoghue. Reach and grasp by people with tetraplegia using a neurally controlled robotic arm. Nature, 485(7398):372–375, 2012.

[17] Jennifer L Collinger, Brian Wodlinger, John E Downey, Wei Wang, Elizabeth C Tyler-Kabara, Douglas J Weber, Angus JC McMorland, Meel Velliste, Michael L Boninger, and Andrew B Schwartz. High-performance neuroprosthetic control by an individual with tetraplegia. The Lancet, 381(9866):557–564, 2013.

[18] Brian Wodlinger, JE Downey, Elizabeth C Tyler-Kabara, Andrew B Schwartz, ML Boninger, and Jennifer L Collinger. Ten-dimensional anthropomorphic arm control in a human brain-machine interface: difficulties, solutions, and limitations. Journal of neural engineering, 12(1):016011, 2015.

[19] Tyson Aflalo, Spencer Kellis, Christian Klaes, Brian Lee, Ying Shi, Kelsie Pejsa, Kathleen Shanfield, Stephanie Hayes-Jackson, Mindy Aisen, Christi Heck, Charles Liu, and Richard A. Andersen. Decoding motor imagery from the posterior parietal cortex of a tetraplegic human. Science, 348(6237):906–910, 2015.

[20] John E Downey, Lucas Brane, Robert A Gaunt, Elizabeth C Tyler-Kabara, Michael L Boninger, and Jennifer L Collinger. Motor cortical activity changes during neuroprosthetic-controlled object interaction. Scientific reports, 7(1):16947, 2017.

[21] Sharlene N Flesher, John E Downey, Jeffrey M Weiss, Christopher L Hughes, Angelica J Herrera, Elizabeth C Tyler-Kabara, Michael L Boninger, Jennifer L Collinger, and Robert A Gaunt. A brain-computer interface that evokes tactile sensations improves robotic arm control. Science, 372(6544):831–836, 2021.

[22] David A. Handelman, Luke E. Osborn, Tessy M. Thomas, Andrew R. Badger, Margaret Thompson, Robert W. Nickl, Manuel A. Anaya, Jared M. Wormley, Gabriela L. Cantarero, David McMullen, Nathan E. Crone, Brock Wester, Pablo A. Celnik, Matthew S. Fifer, and Francesco V. Tenore. Shared control of bimanual robotic limbs with a brain-machine interface for self-feeding. Frontiers in Neurorobotics, 16:918001, 2022.

[23] Syed Abu Huraira Hussain, Imran Raza, Syed Asad Hussain, Muhammad Hasan Jamal, Tauseef Gulrez, and Ali Zia. A mental state aware brain computer interface for adaptive control of electric powered wheelchair. Scientific Reports, 15(1):9880, 2025.

[24] Stephen Rainey, Hannah Maslen, and Julian Savulescu. When thinking is doing: responsibility for BCI-mediated action. AJOB neuroscience, 11(1):46–58, 2020.

[25] Jakob Gawlikowski, Cedrique Rovile Njieutcheu Tassi, Mohsin Ali, Jongseok Lee, Matthias Humt, Jianxiang Feng, Anna Kruspe, Rudolph Triebel, Peter Jung, Ribana Roscher, Muhammad Shahzad, Wen Yang, Richard Bamler, and Xiao Xiang Zhu. A survey of uncertainty in deep neural networks. Artificial Intelligence Review, 56(Suppl 1):1513–1589, 2023.

[26] Sebastian Thrun. Probabilistic robotics. Communications of the ACM, 45(3):52–57, 2002.

[27] Abdullah A Abdullah, Masoud M Hassan, and Yaseen T Mustafa. A review on bayesian deep learning in healthcare: Applications and challenges. IEEe Access, 10:36538–36562, 2022.

[28] Arash Ajoudani, Andrea Maria Zanchettin, Serena Ivaldi, Alin Albu-Schäffer, Kazuhiro Kosuge, and Oussama Khatib. Progress and prospects of the human–robot collaboration. Autonomous robots, 42(5):957–975, 2018.

[29] Reza Abiri, Ali Rabiee, Sima Ghafoori, and Anna Cetera. Toward human-centered shared autonomy ai paradigms for human-robot teaming in healthcare. arXiv preprint 2407.17464, 2024.

[30] Dario Amodei, Chris Olah, Jacob Steinhardt, Paul Christiano, John Schulman, and Dan Mané. Concrete problems in ai safety. arXiv preprint 1606.06565, 2016.

[31] MH Farhadi, Ali Rabiee, Sima Ghafoori, Anna Cetera, Wei Xu, and Reza Abiri. Human-centered shared autonomy for motor planning, learning, and control applications. Bridging the Gap between Mind and Machine: Exploring the Future of Human-AI-Neurotechnology Integration, pages 259– 297, 2025.

[32] Lorenzo Tonin, Tom Carlson, Robert Leeb, and Josédel R. Millán. Combining brain–computer interfaces and assistive technologies: state-of-the-art and challenges. Frontiers in Neuroscience, 9:1–17, 2015.

[33] Christian Herff, Dominic Heger, Adriana De Pesters, Dominic Telaar, Peter Brunner, Gerwin Schalk, and Tanja Schultz. Brain-to-text: decoding spoken phrases from phone representations in the brain. Frontiers in neuroscience, 8:141498, 2015.

[34] Christian Herff and Tanja Schultz. Automatic speech recognition from neural signals: a focused review. Frontiers in neuroscience, 10:429, 2016.

[35] Nicholas S. Card, Maitreyee Wairagkar, Carrina Iacobacci, Xianda Hou, Tyler Singer-Clark, Francis R. Willett, Erin M. Kunz, Chaofei Fan, Maryam Vahdati Nia, Darrel R. Deo, Aparna Srinivasan, Eun Young Choi, Matthew F. Glasser, Leigh R. Hochberg, Jaimie M. Henderson, Kiarash Shahlaie, Sergey D. Stavisky, and David M. Brandman. An accurate and rapidly calibrating speech neuroprosthesis. New England Journal of Medicine, 391(7):609–618, 2024.

[36] Jaimie J. Jude, Hila Levi-Aharoni, A. J. Acosta, S. B. Allcroft, C. Nicolas, B. E. Lacayo, N. S. Card, M. Wairagkar, D. M. Brandman, S. D. Stavisky, F. R. Willett, Z. M. Williams, J. D. Simeral, L. R. Hochberg, and D. B. Rubin. Restoring rapid natural bimanual typing with a neuroprosthesis after paralysis. Nature Neuroscience, 2026.

[37] Sean L. Metzger, Kaylo T. Littlejohn, Alexander B. Silva, David A. Moses, Margaret P. Seaton, Ran Wang, Maximilian E. Dougherty, Jessie R. Liu, Peter Wu, Michael A. Berger, Inga Zhuravleva, Adelyn Tu-Chan, Karunesh Ganguly, Gopala K. Anumanchipalli, and Edward F. Chang. A high-performance neuroprosthesis for speech decoding and avatar control. Nature, 620(7976):1037– 1046, 2023.

[38] Erin M. Kunz, Benyamin Abramovich Krasa, Foram Kamdar, Donald T. Avansino, Nick Hahn, Seonghyun Yoon, Akansha Singh, Samuel R. Nason-Tomaszewski, Nicholas S. Card, Justin J. Jude, Brandon G. Jacques, Payton H. Bechefsky, Carrina Iacobacci, Leigh R. Hochberg, Daniel B. Rubin, Ziv M. Williams, David M. Brandman, Sergey D. Stavisky, Nicholas AuYong, Chethan Pandarinath, Shaul Druckmann, Jaimie M. Henderson, and Francis R. Willett. Inner speech in motor cortex and implications for speech neuroprostheses. Cell, 188(17):4658–4673, 2025.

[39] Samuel R. Nason-Tomaszewski, Pranav I. Deevi, Qinwan Rabbani, Brandon G. Jacques, Anna L. Pritchard, Lahiru N. Wimalasena, Brice A. Richards, Brianna M. Karpowicz, Payton H. Bechefsky, Nicholas S. Card, Darrel R. Deo, Eun Young Choi, Leigh R. Hochberg, Sergey D. Stavisky, David M. Brandman, Nicholas AuYong, and Chethan Pandarinath. Restoring brain-to-text communication in a person with dysarthria from pontine stroke using an intracortical brain-computer interface. medRxiv, pages 2026–02, 2026.

[40] Mehryar Mohri, Fernando Pereira, and Michael Riley. Weighted finite-state transducers in speech recognition. Computer Speech & Language, 16(1):69–88, 2002.

[41] Mehryar Mohri and Michael Riley. Speech recognition with weighted finite-state transducers. In Springer Handbook of Speech Processing, pages 559–584. Springer, 2008.

[42] Yajie Miao, Mohammad Gowayyed, and Florian Metze. EESEN: End-to-end speech recognition using deep RNN models and WFST-based decoding. In 2015 IEEE workshop on automatic speech recognition and understanding (ASRU), pages 167–174. IEEE, 2015.

[43] Alec Radford, Jeffrey Wu, Rewon Child, David Luan, Dario Amodei, and Ilya Sutskever. Language models are unsupervised multitask learners. OpenAI blog, 1(8):9, 2019.

[44] Susan Zhang, Stephen Roller, Naman Goyal, Mikel Artetxe, Moya Chen, Shuohui Chen, Christopher Dewan, Mona Diab, Xian Li, Xi Victoria Lin, Todor Mihaylov, Myle Ott, Sam Shleifer, Kurt Shuster, Daniel Simig, Punit Singh Koura, Anjali Sridhar, Tianlu Wang, and Luke Zettlemoyer. OPT: Open pre-trained transformer language models. arXiv preprint 2205.01068, 2022.

[45] Hugo Touvron, Louis Martin, Kevin Stone, Peter Albert, Amjad Almahairi, Yasmine Babaei, Nikolay Bashlykov, Soumya Batra, Prajjwal Bhargava, Shruti Bhosale, et al. Llama 2: Open foundation and fine-tuned chat models. arXiv preprint 2307.09288, 2023.

[46] Jingyuan Li, Trung Le, Chaofei Fan, Mingfei Chen, and Eli Shlizerman. Brain-to-text decoding with context-aware neural representations and large language models. Journal of Neural Engineering, 22(5):056026, 2025.

[47] Yizi Zhang, Linyang He, Chaofei Fan, Tingkai Liu, Han Yu, Trung Le, Jingyuan Li, Scott Linderman, Lea Duncker, Francis R Willett, Nima Mesgarani, and Liam Paninski. Decoding inner speech with an end-to-end brain-to-text neural interface. arXiv preprint 2511.21740, 2025.

[48] Sheng Feng, Heyang Liu, Yu Wang, and Yanfeng Wang. Towards an end-to-end framework for invasive brain signal decoding with large language models. arXiv preprint 2406.11568, 2024.

[49] Jarod Lévy, Mingfang Zhang, Svetlana Pinet, Jérémy Rapin, Hubert Banville, Stéphane d’Ascoli, and Jean-Rémi King. Brain-to-text decoding: A non-invasive approach via typing. arXiv preprint 2502.17480, 2025.

[50] Johannes Y Lee, Sangjoon Lee, Abhishek Mishra, Xu Yan, Brandon McMahan, Brent Gaisford, Charles Kobashigawa, Mike Qu, Chang Xie, and Jonathan C Kao. Brain–computer interface control with artificial intelligence copilots. Nature Machine Intelligence, pages 1–14, 2025.

[51] Katharina Muelling, Arun Venkatraman, Justin-Stefan Valois, Joseph E. Downey, Jonathan Weiss, Shervin Javdani, Martial Hebert, Andrew B. Schwartz, Jennifer L. Collinger, J. Andrew Bagnell, and Siddhartha S. Srinivasa. Autonomy infused teleoperation with application to brain–computer interface controlled manipulation. Autonomous Robots, 41(6):1401–1422, 2017.

[52] Justin J Jude, Stephanie Haro, Hadar Levi-Aharoni, Hiroaki Hashimoto, Alexander J Acosta, Nicholas S Card, Maitreyee Wairagkar, David M Brandman, Sergey D Stavisky, Ziv M Williams, Sydney S. Cash, John D. Simeral, Leigh R. Hochberg, and Daniel B. Rubin. Decoding intended speech with an intracortical brain-computer interface in a person with longstanding anarthria and locked-in syndrome. bioRxiv, pages 2025–08, 2025.

[53] Ebrahim Feghhi, Shreyas Kaasyap, Nima Ryan Hadidi, and Jonathan C. Kao. Time-masked transformers with lightweight test-time adaptation for neural speech decoding. In Proceedings of the Conference on Neural Information Processing Systems (NeurIPS), 2025. Poster.

[54] Maggie A Collier, Rithika Narayan, and Henny Admoni. The sense of agency in assistive robotics using shared autonomy. In 2025 20th ACM/IEEE International Conference on Human-Robot Interaction (HRI), pages 880–888. IEEE, 2025.

[55] Shervin Javdani, Siddhartha S. Srinivasa, and J. Andrew Bagnell. Shared autonomy via hindsight optimization. In Robotics: Science and Systems (RSS), 2015.

[56] Siddarth Jain and Brenna Argall. Recursive bayesian human intent recognition in shared-control robotics. In 2018 IEEE/RSJ International Conference on Intelligent Robots and Systems (IROS), pages 3902–3909. IEEE, 2018.

[57] Francis R Willett, Jingyuan Li, Trung Le, Chaofei Fan, Mingfei Chen, Eli Shlizerman, Yue Chen, Xin Zheng, Tatsuo S Okubo, Tyler Benster, Hyun Dong Lee, Maxwell Kounga, E. Kelly Buchanan, David Zoltowski, Scott W. Linderman, and Jaimie M. Henderson. Brain-to-text benchmark’24: Lessons learned. arXiv preprint 2412.17227, 2024.

[58] Tyler Benster, Guy Wilson, Reshef Elisha, Francis R Willett, and Shaul Druckmann. A cross-modal approach to silent speech with llm-enhanced recognition. arXiv preprint 2403.05583, 2024.

[59] Jerry Tang, Amanda LeBel, Shailee Jain, and Alexander G Huth. Semantic reconstruction of continuous language from non-invasive brain recordings. Nature Neuroscience, 26(5):858–866, 2023.

[60] Stéphane d’Ascoli, Corentin Bel, Jérémy Rapin, Hubert Banville, Yohann Benchetrit, Christophe Pallier, and Jean-Rémi King. Towards decoding individual words from non-invasive brain recordings. Nature Communications, 16(1):10521, 2025.

[61] Ziyi Ye, Qingyao Ai, Yiqun Liu, Maarten de Rijke, Min Zhang, Christina Lioma, and Tuukka Ruotsalo. Generative language reconstruction from brain recordings. Communications Biology, 8(1):346, 2025.

[62] Zhenhailong Wang and Heng Ji. Open vocabulary electroencephalography-to-text decoding and zero-shot sentiment classification. In Proceedings of the AAAI Conference on Artificial Intelligence, volume 36, pages 5350–5358, 2022.

[63] Chuan Guo, Geoff Pleiss, Yu Sun, and Kilian Q Weinberger. On calibration of modern neural networks. In International conference on machine learning, pages 1321–1330. PMLR, 2017.

[64] Meelis Kull, Telmo M Silva Filho, and Peter Flach. Beyond sigmoids: How to obtain well-calibrated probabilities from binary classifiers with beta calibration. 2017.

[65] Meelis Kull, Miquel Perello Nieto, Markus Kängsepp, Telmo Silva Filho, Hao Song, and Peter Flach. Beyond temperature scaling: Obtaining well-calibrated multi-class probabilities with dirichlet calibration. Advances in neural information processing systems, 32, 2019.

[66] Philipp Röchner, Henrique O Marques, Ricardo JGB Campello, and Arthur Zimek. Evaluating outlier probabilities: assessing sharpness, refinement, and calibration using stratified and weighted measures. Data Mining and Knowledge Discovery, 38(6):3719–3757, 2024.

[67] Glenn Shafer and Vladimir Vovk. A tutorial on conformal prediction. Journal of machine learning research, 9(3), 2008.

[68] Yaniv Romano, Matteo Sesia, and Emmanuel Candes. Classification with valid and adaptive coverage. Advances in neural information processing systems, 33:3581–3591, 2020.

[69] Ranganath Krishnan, Mahesh Subedar, and Omesh Tickoo. Bar: Bayesian activity recognition using variational inference. arXiv preprint 1811.03305, 2018.

[70] Balaji Lakshminarayanan, Alexander Pritzel, and Charles Blundell. Simple and scalable predictive uncertainty estimation using deep ensembles. Advances in neural information processing systems, 30, 2017.

[71] Yarin Gal and Zoubin Ghahramani. Dropout as a bayesian approximation: Representing model uncertainty in deep learning. In international conference on machine learning, pages 1050–1059. PMLR, 2016.

[72] Tilmann Gneiting and Adrian E Raftery. Strictly proper scoring rules, prediction, and estimation. Journal of the American statistical Association, 102(477):359–378, 2007.

[73] Alex Graves. Connectionist temporal classification. In Supervised sequence labelling with recurrent neural networks, pages 61–93. Springer, 2012.

[74] Albert Zeyer, Ralf Schlüter, and Hermann Ney. Why does ctc result in peaky behavior? arXiv preprint 2105.14849, 2021.

[75] Ruizhe Huang, Xiaohui Zhang, Zhaoheng Ni, Li Sun, Moto Hira, Jeff Hwang, Vimal Manohar, Vineel Pratap, Matthew Wiesner, Shinji Watanabe, Daniel Povey, and Sanjeev Khudanpur. Less peaky and more accurate ctc forced alignment by label priors. In ICASSP 2024-2024 IEEE International Conference on Acoustics, Speech and Signal Processing (ICASSP), pages 11831– 11835. IEEE, 2024.

[76] Steve Young, Gunnar Evermann, Mark Gales, Thomas Hain, Dan Kershaw, Xunying Liu, Gareth Moore, Julian Odell, Dave Ollason, Dan Povey, Valtcho Valtchev, and Phil Woodland. The HTK book. Cambridge university engineering department, 3(175):12, 2002.

[77] Hongzhu Li and Weiqiang Wang. Reinterpreting ctc training as iterative fitting. Pattern Recognition, 105:107392, 2020.

[78] Geoffrey E Hinton. Training products of experts by minimizing contrastive divergence. Neural computation, 14(8):1771–1800, 2002.

[79] Meelis Kull and Peter Flach. Novel decompositions of proper scoring rules for classification: Score adjustment as precursor to calibration. In Joint European Conference on Machine Learning and Knowledge Discovery in Databases, pages 68–85. Springer, 2015.

[80] Antonio Bella, Cesar Ferri, José Hernández-Orallo, and María José Ramírez-Quintana. On the effect of calibration in classifier combination. Applied intelligence, 38(4):566–585, 2013.

[81] Morris H DeGroot and Stephen E Fienberg. The comparison and evaluation of forecasters. Journal of the Royal Statistical Society: Series D (The Statistician), 32(1-2):12–22, 1983.

[82] Allan H Murphy. A new vector partition of the probability score. Journal of Applied Meteorology and Climatology, 12(4):595–600, 1973.

[83] Jingya Huang, Aashish N Patel, Sowmya Manojna Narasimha, Gal Mishne, and Vikash Gilja. Word-level error analysis in decoding systems: From speech recognition to brain-computer interfaces. In Interspeech, 2025.

[84] Volodymyr Kuleshov and Percy S Liang. Calibrated structured prediction. Advances in Neural Information Processing Systems, 28, 2015.

[85] Abhyuday Jagannatha and Hong Yu. Calibrating structured output predictors for natural language processing. In Proceedings of the conference. Association for Computational Linguistics. Meeting, volume 2020, page 2078, 2020.

[86] Aviral Kumar and Sunita Sarawagi. Calibration of encoder decoder models for neural machine translation. arXiv preprint 1903.00802, 2019.

[87] Oscar Chang, Dongseong Hwang, and Olivier Siohan. Revisiting the entropy semiring for neural speech recognition. In Proceedings of the International Conference on Learning Representations (ICLR), 2023. Poster.

[88] Tomer Wullach and Shlomo E Chazan. Don’t be so sure! boosting ASR decoding via confidence relaxation. In Proceedings of the AAAI Conference on Artificial Intelligence, volume 37, pages 13780–13788, 2023.

[89] Dong Yu, Jinyu Li, and Li Deng. Calibration of confidence measures in speech recognition. IEEE Transactions on Audio, Speech, and Language Processing, 19(8):2461–2473, 2011.

[90] Hu Liu, Sheng Jin, and Changshui Zhang. Connectionist temporal classification with maximum entropy regularization. Advances in Neural Information Processing Systems, 31, 2018.

[91] John P Cunningham and Byron M Yu. Dimensionality reduction for large-scale neural recordings. Nature neuroscience, 17(11):1500–1509, 2014.

[92] Saurabh Vyas, Matthew D Golub, David Sussillo, and Krishna V Shenoy. Computation through neural population dynamics. Annual Review of Neuroscience, 43(1):249–275, 2020.

[93] Jonathan C Kao, Paul Nuyujukian, Stephen I Ryu, Mark M Churchland, John P Cunningham, and Krishna V Shenoy. Single-trial dynamics of motor cortex and their applications to brain-machine interfaces. Nature Communications, 6:7759, 2015.

[94] Afsheen Afshar, Gopal Santhanam, M Yu Byron, Stephen I Ryu, Maneesh Sahani, and Krishna V Shenoy. Single-trial neural correlates of arm movement preparation. Neuron, 71(3):555–564, 2011.

[95] Amy L Orsborn, Helene G Moorman, Simon A Overduin, Maryam M Shanechi, Dragan F Dimitrov, and Jose M Carmena. Closed-loop decoder adaptation shapes neural plasticity for skillful neuroprosthetic control. Neuron, 82(6):1380–1393, 2014.

[96] Matthew D Golub, Patrick T Sadtler, Emily R Oby, Kristin M Quick, Stephen I Ryu, Elizabeth C Tyler-Kabara, Aaron P Batista, Steven M Chase, and Byron M Yu. Learning by neural reassociation. Nature neuroscience, 21(4):607–616, 2018.

[97] Matthew D Golub, Steven M Chase, Aaron P Batista, and Byron M Yu. Brain–computer interfaces for dissecting cognitive processes underlying sensorimotor control. Current opinion in neurobiology, 37:53–58, 2016.

[98] Matthew D Golub, Byron M Yu, and Steven M Chase. Internal models for interpreting neural population activity during sensorimotor control. Elife, 4:e10015, 2015.

[99] Aashish N Patel, Geeling Chau, Cheng Chang, Allan Sun, Jingya Huang, Tzyy-Ping Jung, and Vikash Gilja. Affective response to volitional input perturbations in obstacle avoidance and target tracking games. In 2021 43rd Annual International Conference of the IEEE Engineering in Medicine & Biology Society (EMBC), pages 6679–6682. IEEE, 2021.

[100] Andrew Myrden and Tom Chau. Effects of user mental state on EEG-BCI performance. Frontiers in human neuroscience, 9:308, 2015.

[101] Songwei Li, Junyi Duan, Yu Sun, Xinjun Sheng, Xiangyang Zhu, and Jianjun Meng. Exploring fatigue effects on performance variation of intensive brain–computer interface practice. Frontiers in Neuroscience, 15:773790, 2021.

[102] Francis R Willett, Daniel R Young, Brian A Murphy, William D Memberg, Christine H Blabe, Chethan Pandarinath, Sergey D Stavisky, Paymon Rezaii, Jad Saab, Benjamin L Walter, Jennifer A. Sweet, Jonathan P. Miller, Jaimie M. Henderson, Krishna V. Shenoy, John D. Simeral, Beata Jarosiewicz, Leigh R. Hochberg, Robert F. Kirsch, and A. Bolu Ajiboye. Principled BCI decoder design and parameter selection using a feedback control model. Scientific reports, 9(1):8881, 2019.

[103] Francis R Willett, Chethan Pandarinath, Beata Jarosiewicz, Brian A Murphy, William D Memberg, Christine H Blabe, Jad Saab, Benjamin L Walter, Jennifer A Sweet, Jonathan P Miller, Jaimie M Henderson, Krishna V Shenoy, John D Simeral, Leigh R Hochberg, Robert F Kirsch, and A Bolu Ajiboye. Feedback control policies employed by people using intracortical brain–computer interfaces. Journal of Neural Engineering, 14(1):016001, 2017.

[104] Gakuto Kurata and Kartik Audhkhasi. Guiding ctc posterior spike timings for improved posterior fusion and knowledge distillation. arXiv preprint 1904.08311, 2019.

[105] Matthew S Willsey, Nishal P Shah, Donald T Avansino, Nick V Hahn, Ryan M Jamiolkowski, Foram B Kamdar, Leigh R Hochberg, Frangis R Willett, and Jaimie M Henderson. A highperformance brain–computer interface for finger decoding and quadcopter game control in an individual with paralysis. Nature Medicine, 31(1):96–104, 2025.

[106] Nishal P Shah, Donald Avansino, Foram Kamdar, Claire Nicolas, Anastasia Kapitonava, Carlos Vargas-Irwin, Leigh R Hochberg, Chethan Pandarinath, Krishna V Shenoy, Francis R Willett, and Jamie H Henderson. Pseudo-linear summation explains neural geometry of multi-finger movements in human premotor cortex. Nature Communications, 16(1):5008, 2025.

[107] Matthew S Willsey, Samuel R Nason-Tomaszewski, Scott R Ensel, Hisham Temmar, Matthew J Mender, Joseph T Costello, Parag G Patil, and Cynthia A Chestek. Real-time brain-machine inter-face in non-human primates achieves high-velocity prosthetic finger movements using a shallow feedforward neural network decoder. Nature Communications, 13(1):6899, 2022.

[108] Maneeshika M Madduri, Momona Yamagami, Si Jia Li, Sasha Burckhardt, Samuel A Burden, and Amy L Orsborn. Computational framework to predict and shape human–machine interactions in closed-loop, co-adaptive neural interfaces. Nature Machine Intelligence, pages 1–16, 2026.

[109] Karunesh Ganguly and Jose M Carmena. Emergence of a stable cortical map for neuroprosthetic control. PLoS biology, 7(7):e1000153, 2009.

[110] Andrew Jackson and Eberhard E Fetz. Interfacing with the computational brain. IEEE Transactions on Neural Systems and Rehabilitation Engineering, 19(5):534–541, 2011.

[111] Emily R Oby, Matthew D Golub, Jay A Hennig, Alan D Degenhart, Elizabeth C Tyler-Kabara, Byron M Yu, Steven M Chase, and Aaron P Batista. New neural activity patterns emerge with long-term learning. Proceedings of the National Academy of Sciences, 116(30):15210–15215, 2019.

